# Quantitative modeling of the emergence of macroscopic grid-like representations

**DOI:** 10.1101/2022.12.20.521210

**Authors:** Ikhwan Bin Khalid, Eric T. Reifenstein, Naomi Auer, Lukas Kunz, Richard Kempter

## Abstract

Grid cells are neurons in the entorhinal cortex that are thought to perform neural computations in support of spatial navigation. When subjects navigate through spatial environments, grid cells exhibit firing fields that are arranged in a triangular grid pattern. As direct recordings of grid cells from the human brain are only rarely possible, functional magnetic resonance imaging (fMRI) studies proposed and described an indirect measure of entorhinal grid-cell activity, which is quantified as a hexadirectional modulation of fMRI activity as a function of the subject’s movement direction. However, it still remains unclear how the activity of a population of grid cells may exhibit hexadirectional modulation and thus provides the basis for the hexadirectional modulation of entorhinal cortex activity measured with fMRI. Here, we thus performed numerical simulations and analytical calculations to better understand how the aggregated activity of many grid cells may be hexadirectionally modulated. Our simulations implemented three different hypotheses proposing that the hexadirectional modulation occurs because grid cells show head-direction tuning aligned with the grid axes; are subjected to repetition suppression; or exhibit a bias towards a particular grid phase offset. Our simulations suggest that hexadirectional modulation is best explained by the conjunctive grid by head-direction cell hypothesis, which can produce the strongest and most robust hexasymmetry. In contrast, our simulations including previously observed biological properties of grid cells do not provide clear support for the structure-function mapping hypothesis. Our observations on hexadirectional modulation generated by grid-cell adaptation effects and the available data on adaptation properties of grid cells are insufficient to substantiate or refute the repetition suppression hypothesis. Furthermore, we found that the magnitude of the hexadirectional modulation depends considerably on the subject’s navigation pattern. Our results thus indicate that future fMRI studies could be designed to test which of the three hypotheses most likely accounts for the fMRI measure of grid cells. These findings also underline the importance of quantifying the biological properties of single grid cells in humans to further elucidate how hexadirectional modulations of fMRI activity may emerge.

## 1 Introduction

The neural basis of spatial navigation comprises multiple specialized cell types such as place cells (O’Keefe and Dostrovsky, 1971), head-direction cells (Taube et al., 1990), and grid cells (Hafting et al., 2005), whose activity profiles result from intricate mechanisms of microcircuits in the medial temporal lobes (Tukker et al., 2022). Grid cells are neurons that activate whenever an animal or human traverses the vertices of a triangular grid tiling the entire environment into equilateral triangles (Hafting et al., 2005; Jacobs et al., 2013). Grid cells may allow the navigating organism to perform vector computations and may thus constitute an essential neural substrate for different types of spatial navigation including path integration, though their exact functional role still remains unclear (Stemmler et al., 2015; Bush et al., 2015; Moser et al., 2017; Stangl et al., 2018; Gil et al., 2018; Banino et al., 2018; Bierbrauer et al., 2020; Ginosar et al., 2023).

In rodents, grid cells can be recorded using electrodes inserted into the medial entorhinal cortex (EC). In humans, measuring grid cells using invasive methods is only rarely possible, for example, by recording single-neuron activity in epilepsy patients who are neurosurgically implanted with intracranial depth electrodes (Jacobs et al., 2013; Nadasdy et al., 2017). Hence, to enable the detection of grid cells in healthy humans, a functional magnetic resonance imaging (fMRI) method has been developed that tests for a hexadirectional modulation of the blood-oxygen-level-dependent (BOLD) signal as a function of the subject’s movement direction through a virtual environment (Doeller et al., 2010). We here refer to this phenomenon of a hexadirectional modulation of the fMRI signal as “macroscopic grid-like representations”, which has been replicated repeatedly in recent years (e.g. Kunz et al., 2015; Bellmund et al., 2016; Horner et al., 2016; Constantinescu et al., 2016; Bierbrauer et al., 2020; Convertino et al., 2023). The mechanisms underlying the emergence of such macroscopic grid-like representations remain still unclear, however.

To provide possible explanations for the emergence of macroscopic grid-like representations, previous studies presented several qualitatively different hypotheses on how the activity of single grid cells translates into a macroscopically visible hexadirectional fMRI signal (Doeller et al., 2010; Kunz et al., 2019). Three main hypotheses have been developed: (i) the “conjunctive grid by head-direction cell hypothesis”; (ii) the “repetition suppression hypothesis”; and (iii) the “structure-function mapping hypothesis” (Fig. 1).

**Figure 1:**
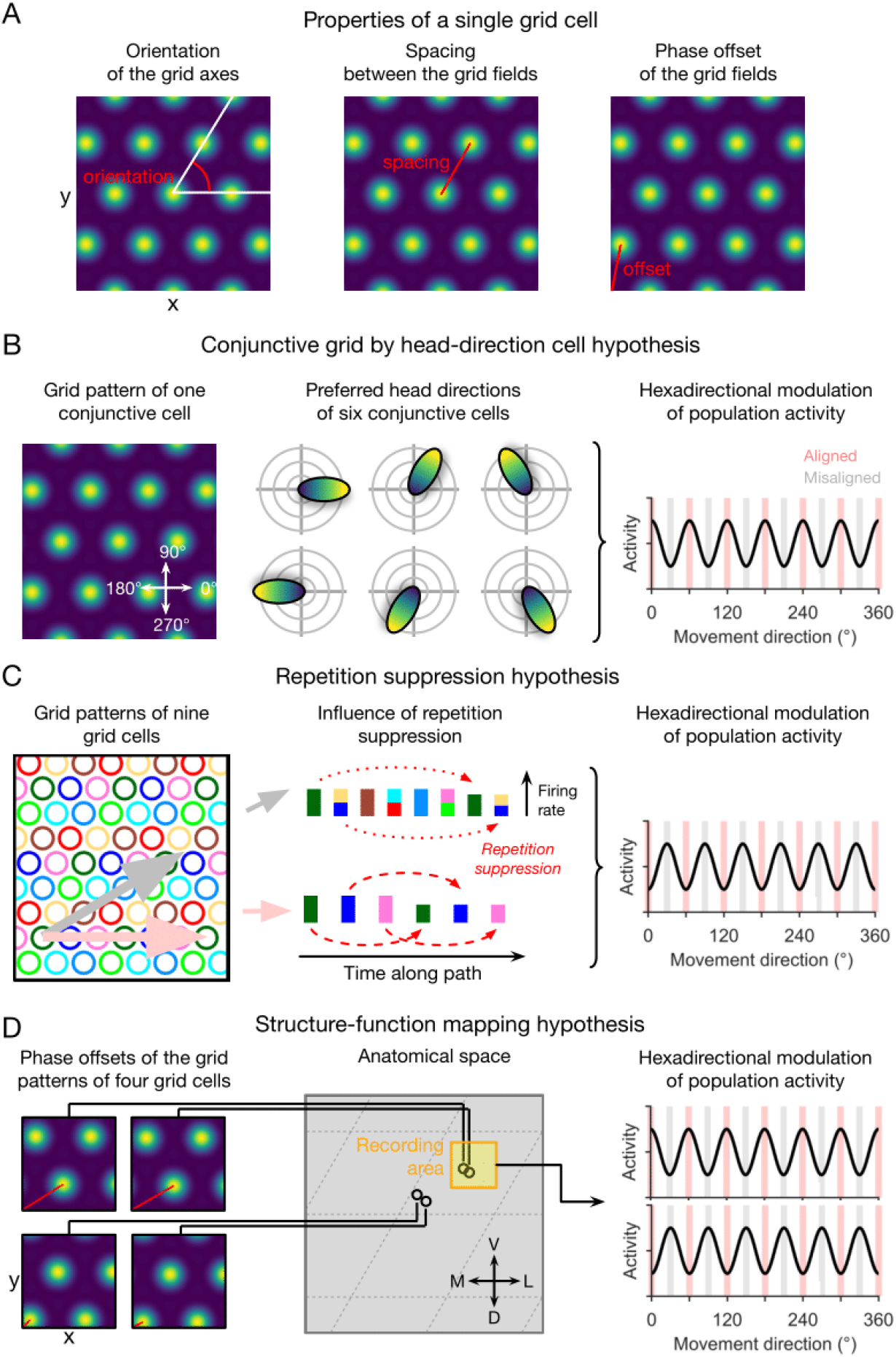
Qualitative hypotheses on the emergence of macroscopic grid-like representations in the human entorhinal cortex (adapted from Kunz et al., 2019). (**A**) Grid-cell properties comprise grid orientation, grid spacing, and grid phase offset. (**B**) Conjunctive grid by head-direction cell hypothesis. Macroscopic grid-like representations (right) may emerge from the firing of conjunctive grid by head-direction cells (left and middle) that exhibit increased firing when the subject moves aligned as compared to misaligned with the grid axes (right) (Doeller et al., 2010). (**C**) Repetition suppression hypothesis. In a grid cell population with similar grid orientations and grid spacings but with distributed phases (left; colored circles represent firing fields of different grid cells), aligned movements (horizontal pink arrow) lead to more frequent activation (shorter distance between firing fields) of a smaller number of different grid cells, whereas misaligned movements (diagonal gray arrow) lead to less frequent activation (larger distance between firing fields) of a higher number of different grid cells (Doeller et al., 2010). Thus, aligned movements may lead to more pronounced repetition suppression as compared to misaligned movements (middle), resulting in a hexadirectional modulation of population spiking activity and thus in the emergence of grid-like representations (right). (**D**) Structure–function mapping hypothesis. Because anatomically adjacent grid cells exhibit similar grid phase offsets (in addition to similar grid orientations and grid spacings) (Gu et al., 2018), recordings from a limited number of grid cells with a non-random distribution of phase offsets may lead to macroscopic grid-like representations. The left panel shows the grid phase offset of four different grid cells, whose anatomical locations are illustrated in the middle panel. Depending on the subject’s starting location relative to the phase offset of the grid fields, movements aligned or misaligned with the grid axes lead to higher sum grid cell activity as compared to misaligned or aligned movements (right panel). Furthermore, the orientation of hexadirectional modulation may shift when recording from neighboring voxels in anatomical space due to a shift in the clustered phase offsets. D, dorsal; L, lateral; M, medial; V, ventral.

The conjunctive grid by head-direction cell hypothesis rests on two key findings: first, the existence of conjunctive grid by head-direction cells, which were found in the deeper layers of the entorhinal cortex and in pre- and parasubiculum (Sargolini et al., 2006); second, the observation that the directional tuning of grid cells within the entorhinal cortex is aligned with the grid axes (Doeller et al., 2010), though further studies are needed to corroborate this observation. Assuming that the directional tuning width of these conjunctive grid by head-direction cells is not too broad, movements aligned with the grid axes (as compared to misaligned movements) result in increased spiking activity of the conjunctive grid by head-direction cell population. Given some correlation between population spiking activity and the fMRI signal, this systematic difference in the firing of conjunctive grid by head-direction cells when moving aligned versus misaligned with the grid axes may thus cause a macroscopically visible fMRI signal with hexadirectional modulation (Fig. 1B).

We note that the conjunctive grid by head-direction cell hypothesis does not necessarily depend on a tight alignment between the preferred head directions with the grid axes. As long as the offset between different preferred head directions is at multiples of 60 degrees, any constant offset to the grid axes (not only 0 degrees) will lead to the same hexasymmetry. The conjunctive grid by head-direction cell hypothesis thus also works without grid cells, which may explain why grid-like representations have been observed (using fMRI) in regions outside the entorhinal cortex, where rodent studies have not yet identified grid cells (Doeller et al., 2010; Constantinescu et al., 2016). In that case, however, another mechanism would be needed that could explain why the preferred head directions of different head-direction cells occur at multiples of 60 degrees. Attractor-network structures may be involved in such a mechanism, but this remains speculative at the current stage.

The repetition suppression hypothesis (Fig. 1C) is based on the assumption that the phenomenon of repetition suppression—i.e., neural activity being reduced for repeated stimuli (Grill-Spector et al., 2006)—also applies to grid cells (Doeller et al., 2010; Killian et al., 2012). Critical to this hypothesis is that relatively fewer different grid cells are activated more often during movements aligned with the grid axes, and relatively more different grid cells are activated less often during misaligned movements. Due to this systematic difference in how many grid cells are activated how often, a higher degree of repetition suppression at the level of spiking activity or the fMRI signal (i.e., fMRI adaptation) during aligned movements as compared with misaligned movements can emerge, again resulting in a hexadirectional modulation of fMRI activity as a function of the subject’s movement direction through the spatial environment.

Regarding the structure–function mapping hypothesis (Fig. 1D), studies in rodents have demonstrated that the firing fields of anatomically adjacent grid cells do not only have similar spacing and orientation (Stensola et al., 2012), but also a similar grid phase offset to a reference location (Heys et al., 2014; Gu et al., 2018). Studies in rodents observed that grid cells also cluster anatomically (Obenhaus et al., 2022; Naumann et al., 2018), which may lead to stronger grid-like representations in some voxels (with an increased percentage of grid cells) than in others (with a decreased percentage of grid cells). Therefore, recordings from a small area of the entorhinal cortex (e.g. a sufficiently small voxel of an fMRI scan) may sample grid cells with similar firing fields, which basically behave similarly. It has been suggested that such a grid cell population might show higher average firing rates during aligned movements (because more firing fields are traversed) versus misaligned movements, again resulting in macroscopically visible grid-like representations (Kunz et al., 2019).

In this study, we aimed at quantitatively evaluating the three hypotheses on the emergence of macroscopic grid-like representations using a modeling approach. Our results show that all three hypotheses can result in macroscopic grid-like representations under ideal and specific conditions, but that the magnitude of the hexadirectional modulation varies by orders of magnitude. Key findings are also that the subjects’ type of navigation paths through the spatial environments and the exact biological characteristics of grid cells determine to what extent a given hypothesis can explain a hexadirectional population signal in the entorhinal cortex. In this way, our results help understand how grid cells may have a specific correlate in fMRI, make predictions on how future fMRI studies could establish evidence in favor of or against one of the three hypotheses, and suggest that the biological properties of grid cells in humans should be investigated in greater detail in order to support or weaken the plausibility of either of the three hypotheses.

## 2 Results

In what follows, we first describe the navigation strategies that we use in our model, then define a new measure to quantify neural hexasymmetry and path hexasymmetry, and finally calculate neural hexasymmetries for the three hypotheses, which we model for idealized as well as realistic choices of parameters.

### 2.1 Navigation strategies

To evaluate the different hypotheses on the emergence of grid-like representations, we considered three different types of navigation trajectories: “star-like walks”, “piecewise linear walks”, and “random walks”. We opted for this approach in order to test whether a subject’s navigation pattern—which in itself comes with a certain degree of hexasymmetry (“path hexasymmetry”)—influences the emergence of hexadirectional sum signals of neuronal activity.

During each path segment of star-like walks, the simulated agent started from the same (x/y)-coordinate and navigated along one of 360 predefined allocentric navigation directions (0° to 359° in steps of 1°; Fig. 2B). This ensured that the navigation trajectory itself exhibited a hexasymmetry that was essentially zero. Each path had a length of 300 cm, which was ten times the grid scale (see Table 1 for a summary of all model parameters). After each path segment, the agent was “teleported” back to the initial (x/y)-coordinate and completed the next path segment. For real-world experiments, this type of navigation including teleportation is unusual, but it can be implemented in virtual-reality experiments (Vass et al., 2016; Deuker et al., 2016).

**Figure 2:**
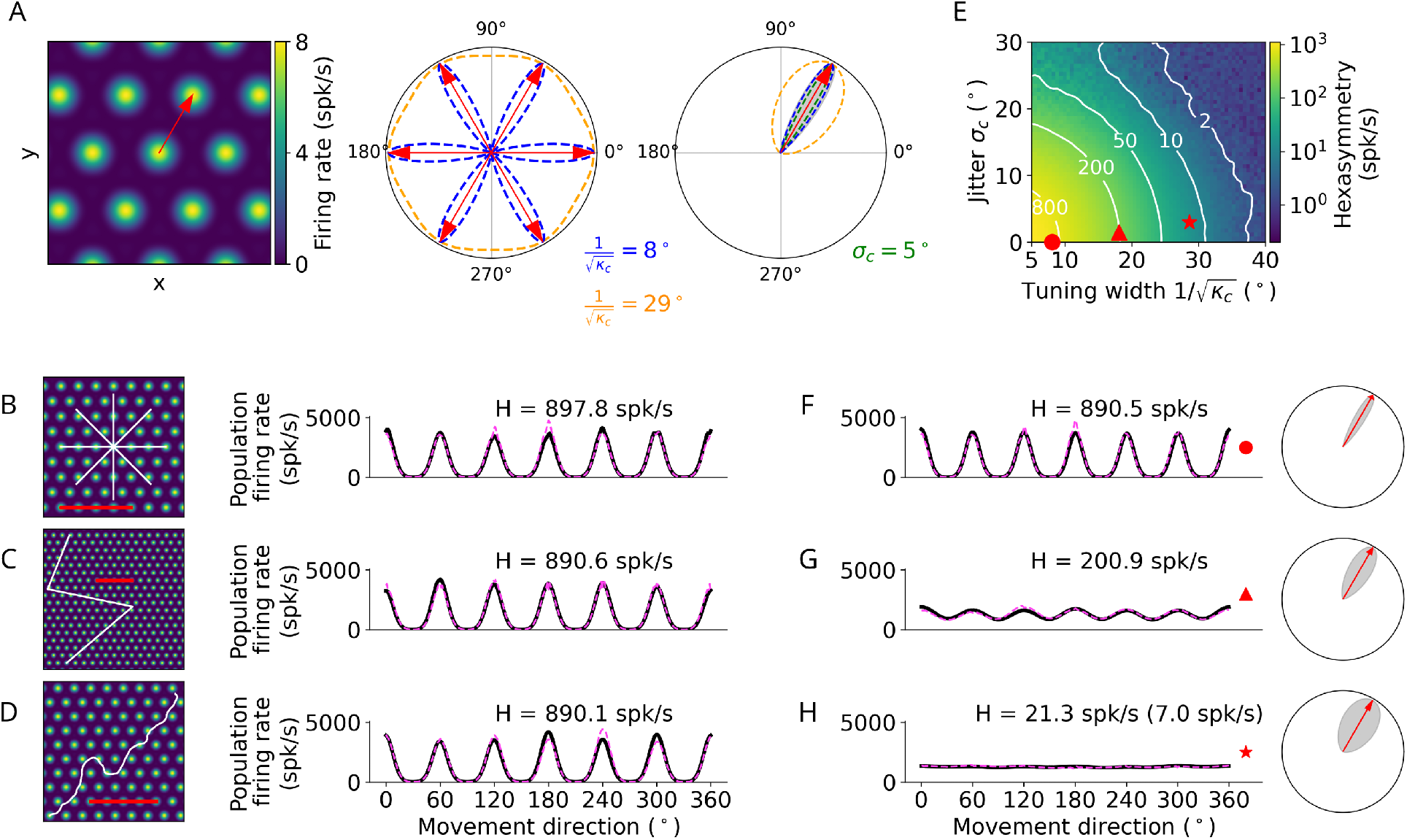
Conjunctive grid by head-direction cell hypothesis. (**A**) Left: The preferred head direction (red arrow) of a single conjunctive cell is aligned to one of the grid axes (Doeller et al., 2010). Two factors add noise to this relation: the HD tuning has a certain width 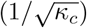 and the alignment of grid orientation to HD tuning angle is jittered (*σ*_*c*_). Middle: Distribution of the tuning width 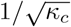 around all possible grid axes for two example values. Right: Convolving the distributions of the jitter *σ*_*c*_ = 5° and tuning width 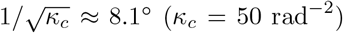 to obtain the effective head-direction (HD) tuning of a single conjunctive cell around one grid axis. (**B–D**) Simulation of the conjunctive hypothesis using “ideal” parameters of *κ*_*c*_ = 50 rad^*−*2^ and *σ*_*c*_ = 0°. The scale bars (red) represent a distance of 120 cm. (**B**) Left: Illustration of a star-like walk (path segments are cut for illustration purposes), overlaid onto the firing pattern of a single grid cell. Right: Population firing rate as a function of the subject’s movement direction (which is identical with heading direction in our simulations) for star-like runs with mean firing rate *Ã*_0_ = 1279.7 spk/s (for 1024 cells) and path hexasymmetry 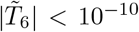 see Methods for definitions of *Ã*_0_ and 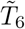. (**C**) Left: Illustration of a piecewise linear walk (cut for illustration purposes), overlaid onto the firing pattern of a single grid cell. Right: Population firing rate as a function of movement direction for piecewise linear walks with *Ã*_0_ = 1279.3 spk/s and 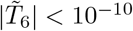 (**D**) Left: Illustration of a random walk (cut for illustration purposes), overlaid onto the firing pattern of a single grid cell. Right: Population firing rate as a function of movement direction for random walks with *Ã*_0_ = 1281.0 spk/s and 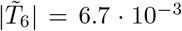. (**E**) Hexasymmetry (color coded) as a function of HD tuning width and alignment jitter for star-like walk trajectories. Higher hexasymmetry values are achieved for stronger HD tuning and tighter alignment of the preferred head directions to the grid axes. The red symbols correspond to the three parameter combinations used in subplots (B–D and F–H) for further illustration. Large tuning widths (*κ*_*c*_*→* 0) correspond to cosine tuning for which the hexasymmetry approaches 0. (**F**) Population firing rate as a function of movement direction for a random walk trajectory with jitter *σ*_*c*_ = 0 and concentration parameter *κ*_*c*_ = 50 rad^*−*2^ (tuning width *≈*8.1°). (**G**) Population firing rate as a function of movement direction for a random walk with jitter *σ*_*c*_ = 1.5° and concentration parameter *κ*_*c*_ = 10 rad^*−*2^ (tuning width *≈*18.1°). (**H**) Population firing rate as a function of movement direction for a random walk with jitter *σ*_*c*_ = 3° and concentration parameter *κ*_*c*_ = 4 rad^*−*2^ (tuning width *≈* 28.6°). The hexasymmetry for the case of *p*_*c*_ = 33% is stated in brackets. All other simulations presented in this figure use *p*_*c*_ = 100% conjunctive (*N* = 1024) cells, which is higher than in empirical studies (Sargolini et al., 2006; Boccara et al., 2010). In subplots B–D and F–H, the black solid lines and magenta dashed lines represent the results from the numerical simulations of Eq. (8) and the analytical derivation in Eq. (32), respectively. The radial subplots in F–H, right, illustrate the effective HD tuning width around a single grid axis analogous to A, right. H, neural hexasymmetry; spk/s, spikes per second.

**Table 1:** Parameters: descriptions and values. For a more detailed motivation of the values used, see section 5.6 Parameter estimation.

| Parameter | Description | Values (unless varied) or Range |
| --- | --- | --- |
| **Trajectories** |  |  |
| $\Delta t$ | Simulation time step | 0.01 s |
| $T$ | Simulated duration | 9000 s |
| $v$ | Movement speed | 10 cm/s |
| $\sigma_\theta$ | Movement tortuosity | $0.5 \text{ rad/s}^{1/2}$ |
| $r_{max}$ | Length of a linear path in the star-like run | 300 cm |
| $N_\theta$ | Number of angles to sample in the star-like run | 360 |
| **Grid cells** |  |  |
| $N$ | Number of grid cells in a voxel | 1024 |
| $s$ | Grid scale | 30 cm |
| $\gamma$ | Grid orientation | $0^\circ$ |
| $A_{max}$ | Maximum firing rate for one grid cell | 8 spk/s |
| $(x_{off}, y_{off})$ | Grid phase (2-dimensional) | $([0,1], [0,1])$ |
| **Conjunctive grid by head-direction cell hypothesis** |  |  |
| $\mu_c$ | Preferred head direction | $[0, 2\pi)$ (multiples of $60^\circ$ for $\sigma_c = 0$) |
| $\kappa_c$ | Concentration parameter for direction tuning for the ideal and “realistic” cases | $\{50, 4\} \text{ rad}^{-2}$ |
| $\sigma_c$ | Alignment jitter of direction tuning to grid axis for the ideal and “realistic”<sup>1</sup> cases | $\{0, 3\}^\circ$ |
| $p_c$ | Fraction of conjunctive cells in a population for the ideal and “realistic”<sup>1</sup> cases | $\{1, 1/3\}$ |
| **Repetition suppression hypothesis** |  |  |
| $\tau_r$ | Adaptation time constant for the ideal and “weaker” cases | $\{3, 1.5\} \text{ s}$ |
| $w_r$ | Adaptation weight for the ideal and “weaker” cases | $\{1, 0.5\}$ |
| **Structure-function mapping hypothesis** |  |  |
| $\mu_s$ | Central phase of cluster | $(0, 0)$ |
| $\kappa_s$ | Concentration parameter for clustering for the ideal and “realistic”<sup>2</sup> cases | $\{10, 0.1\}$ |

<sup>1</sup>(Boccaro et al., 2010; Sargolini et al., 2006)
<sup>2</sup>(Gu et al., 2018)

During piecewise linear walks, the subject also completed 360 path segments of 300 cm length along the same 360 predefined allocentric navigation directions, as in the star-like walks. In this case, however, the path segments were “unwrapped” such that the starting location of a path segment was identical with the end location of the preceding path segment (Fig. 2C). The sequence of allocentric navigation directions was randomly chosen. As for star-like walks, piecewise linear walks do not exhibit hexasymmetry *a priori*.

For random walks, we modeled navigation trajectories following (Kropff and Treves, 2008; Si et al.,2012; D’Albis and Kempter, 2017)(for details, see Methods around Eq. (1)), which allowed us to vary the tortuosity of the paths. For a certain value of the tortuosity parameter (*σ*_*θ*_ = 0.5 rad/s^1/2^) and a time step Δ*t* = 0.01 s, this led to navigation paths that we considered similar to those seen in rodent studies (Fig. 2D; Fig. S2E). Tortuosity describes how convoluted a navigation path is (the higher the tortuosity, the more convoluted the path). A value of 0.5 rad/s^1/2^ means that in one second of time the standard deviation of movement direction change is about 0.5 rad= 29 degrees (see Figure S2E, and Equation 42 for *m*Δ*t* = 1 s). Apart from random walks in basically infinite environments, we also simulated random walks in finite enclosures (circles and squares) with different sizes and orientations; we found that these restrictions had a negligible effect on path hexasymmetry (Figs. S5C and S6C). Because the allocentric navigation directions are not predefined for random walks, they exhibit varying degrees of path hexasymmetry. The longer the simulated random walks, the smaller the path hexasymmetry (Fig. S1). We simulated random walks with a total length of typically 900 m (*M* = 9 · 10^5^ steps).

As we describe below, the emergence of grid-like representations based on the conjunctive hypothesis is robust against the specific type of navigation strategy, whereas the other two hypotheses are sensitive to particular navigation strategies. Future studies on hexadirectional signals should thus consider the kind of navigation paths subjects will use during a given task.

### 2.2 Quantifying neural hexasymmetry generated by the three hypotheses

To test how the activity of grid cells could give rise to hexasymmetry of a macroscopic signal, we used a firing-rate model of grid cell activity (Eq. 2). Furthermore, we developed a new measure *H* to quantify neural hexasymmetry (see Methods, Eq. 12), which is the magnitude of the hexadirectional modulation of the summed activity of many cells. The hexasymmetry *H* has a value *H* = 0 if there is no hexadirectional modulation, i.e., if the population firing rate does not depend on the movement direction. Conversely, if the population firing rate has a hexadirectional sinusoidal modulation, half of its amplitude (in units of the firing rate) equals the value of the hexasymmetry *H*. Using the same approach, we quantified the hexasymmetry of a trajectory and called this path hexasymmetry (see Methods after Eq. 14).

#### 2.2.1 Conjunctive grid by head-direction cell hypothesis

The conjunctive grid by head-direction cell hypothesis (Doeller et al., 2010) suggests that hexadirectional activity in the entorhinal cortex emerges due to grid cells whose firing rate is additionally modulated by head direction, whereby the preferred head direction is closely aligned with one of the grid axes (Fig. 2A). By modulating the activity of individual grid cells with a head-direction tuning term aligned with the grid axes (Methods, Eq. 3), our simulations indeed showed that these properties resulted in a clear hexadirectional modulation of sum grid-cell activity (Fig. 2, B–D). When considering different types of navigation trajectories, we found that they led to similar distributions of sum grid-cell activity as a function of movement direction and, accordingly, to similar hexasymmetry values (Fig. 2, B–D). In all three cases, the directions of maximal activity were aligned with the grid axes.

These results were obtained using ideal values for the preciseness of the head-direction tuning (i.e., the concentration parameter of head-direction tuning, *κ*_*c*_) and the alignment of the preferred head directions to the grid axes (i.e., the alignment jitter of the head-direction tuning to the grid axes, *σ*_*c*_; Fig. 2A). We were thus curious how the hexasymmetry changed when using a wide range of parameter values that would also include biologically plausible values. We varied *κ*_*c*_ between values corresponding to narrow tuning widths (*κ*_*c*_ = 50 rad^*−*2^, which corresponds to an angular variability of 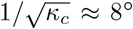) and wide tuning widths (*κ*_*c*_ = 4 rad^*−*2^, i.e. an angular variability of approximately 29°), and *σ*_*c*_ between values of no jitter (*σ*_*c*_ = 0) and significant jitter (*σ*_*c*_ = 3°); see also Table 1 for a summary of parameters. We found that a combination of narrow head direction tuning widths and no jitter resulted in the largest hexasymmetry *H* (Fig. 2, E–H), while wider tuning widths with non-zero jitter resulted in smaller values for the hexasymmetry.

#### 2.2.2 Repetition suppression hypothesis

Next, we performed simulations to understand whether the repetition suppression hypothesis (Doeller et al., 2010) results in a hexadirectional modulation of population grid-cell activity. This hypothesis proposes that grid-cell activity is subject to firing-rate adaptation and thus leads to reduced grid-cell activity when moving along the grid axes as compared to when moving along other directions than the grid axes (Fig. 3A). This difference is due to the fact that the grid fields of fewer grid cells are traversed relatively more often when the subject moves along the grid axes (associated with strong repetition suppression), whereas the grid fields of more grid cells are traversed relatively less often when moving not along the grid axes (weak repetition suppression) (Doeller et al., 2010).

**Figure 3:**
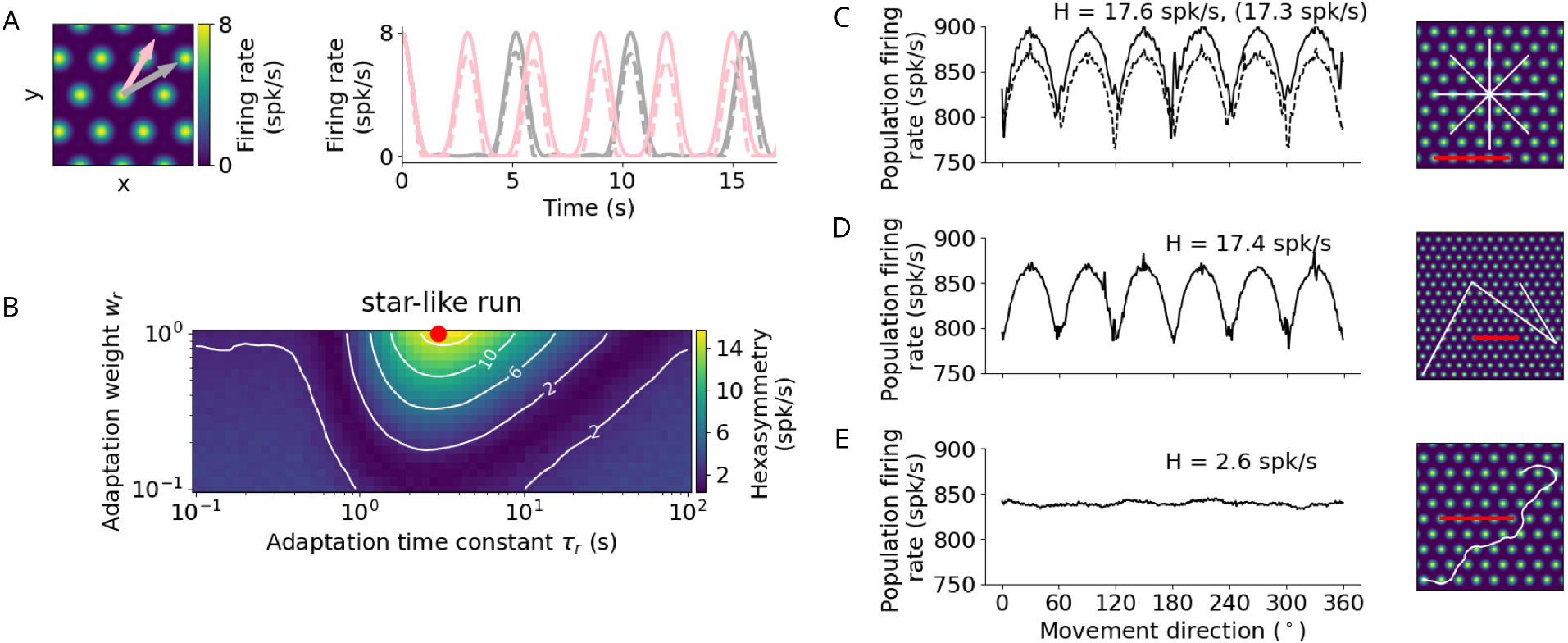
Repetition suppression hypothesis. (**A**) Left: Tuning of an example grid cell with aligned (pink arrow) and misaligned (gray arrow) movement directions. Right: Examples of firing-rate adaptation (dashed lines, adaptation weight *w*_*r*_ = 1 and time constant *τ*_*r*_ = 3 s) for an aligned run (pink) and a misaligned run (gray). More firing-rate adaptation (i.e., stronger repetition suppression) occurs along the aligned run compared to the misaligned run (attenuation of peaks: 24% and 18% respectively). For both runs, firing rates are reduced compared to the case without adaptation (solid lines). (**B**) Simulations of hexasymmetry as a function strength (*w*_*r*_) and time constant (*τ*_*r*_) of adaptation for star-like runs. Red dot marks the optimal parameters (*w*_*r*_ = 1 and *τ*_*r*_ = 3 s) used in (A, C–E). (**C**) Population firing rate as a function of the subject’s movement direction for a star-like run (at an offset of (0, 0)). The solid line represents a single run where adaptation does not carry over when sampling different movement directions (i.e., the “teleportation” between path segments resets the repetition suppression effects), and movement directions are sampled consecutively from 0° to 359° in steps of 1° (mean firing rate *Ã*_0_ = 866.4 spk/s, path hexasymmetry 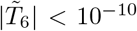). The dashed line represents a single run with adaptation carry-over and randomly sampled movement directions without replacement (*Ã*_0_ = 839.8 spk/s, path hexasymmetry 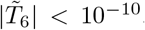, the corresponding hexasymmetry is shown in brackets). (**D**) Population firing rate as a function of movement direction for a piecewise linear walk (*Ã*_0_ = 839.0 spk/s, 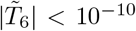). (**E**) Population firing rate as a function of movement direction for a random walk (*Ã*_0_ = 839.7 spk/s, 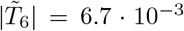). spk/s, spikes per second. For all repetition-suppression simulations, the grid phase offsets of 1024 grid cells were sampled randomly from a uniform distribution across the unit rhombus, and the hexasymmetries were averaged over 20 realisations. The scale bars (red) in (C–E) represent a distance of 120 cm.

In our model, the repetition suppression hypothesis depends on two adaptation parameters: the adaptation time constant *τ*_*r*_ and the adaptation weight *w*_*r*_ (Eq. 5). We explored a large range of adaption time constants and found that the time constant that leads to the largest hexasymmetry is roughly the subject’s speed *v* divided by the grid scale *s* (Fig. 3B). We constrained values of the adaptation weight to the full range of reasonable values (0 < *w*_*r*_ ≤ 1) and found that the larger the value of the adaptation weight the larger is the hexasymmetry (Fig. 3B). When examining how the different types of navigation trajectories affected hexadirectional modulations based on repetition suppression, we found that star-like and piecewise linear walks resulted in clear and significant hexasymmetry values (Fig. 3C, D), which is driven by the long linear segments in these trajectory types. In contrast, random walks did not result in a significant hexadirectional modulation of sum grid-cell activity because the tortuosity of the random walk that we typically used (*σ*_*θ*_ = 0.5 rad/s^1/2^) is too large (i.e. trajectories turn too much between two adjacent firing fields of a grid cell) to be able to exploit the movement-direction dependence of repetition suppression. Examining in more detail which tortuosity values would still lead to some hexadirectional modulation due to repetition suppression, we found that *σ*_*θ*_ ≲ 0.25 rad/s^1/2^ was the upper bound. For smaller values of the tortuosity parameter trajectories are straight enough to allow for a hexadirectional modulation of sum grid-cell activity (Fig. S2).

A notable difference of the repetition suppression hypothesis compared to the conjunctive grid by head-direction cell hypothesis is that the apparent preferred grid orientation (i.e., the movement directions resulting in the highest sum grid-cell activity) is shifted by 30° and is thus exactly misaligned with the grid axes of the individual grid cells (Fig. 3, C–D). This is due to the fact that the adaptation mechanism suppresses grid-cell activity more strongly when moving aligned with a grid axis as compared to when moving misaligned with a grid axis (Fig. 3A).

#### 2.2.3 Structure-function mapping hypothesis

We next investigated the structure-function mapping hypothesis, according to which a hexadirectional modulation of entorhinal cortex activity emerges in situations when a population of grid cells is recorded whose grid phase offsets are biased towards a particular offset (Kunz et al., 2019). In the ideal case, all grid phase offsets are identical (and thus all grid cells behave like a single grid cell). This hypothesis is called “structure-function mapping hypothesis” because of a direct mapping between the anatomical locations of the grid cells in the entorhinal cortex and their functional firing fields in space.

We found indeed that highly clustered grid phase offsets (*κ*_*s*_ = 10) resulted in significant hexadirectional modulations of sum grid-cell activity when the subject performed star-like walks starting at a phase offset of (0, 0), i.e., the center of the cluster of firing fields of grid cells (Fig. 4A). Interestingly, the hexasymmetry values during star-like walks were strongly dependent on the subject’s starting location relative to the locations of the grid fields: only particular starting phases (within the unit rhombus of grid phase offsets) such as (0, 0) or (0.3, 0.3) led to clear hexasymmetry whereas others, e.g. (0.6, 0), did not (Fig. 4D, left). Additionally, the “apparent preferred grid orientation” (i.e., the movement directions associated with the highest sum grid-cell activity) was shifted by 30° for certain offsets in the unit rhombus illustrating the subject’s starting locations relative to the firing-field locations (Fig. 4D, right). It was furthermore notable that the summed grid-cell activity as a function of movement direction exhibited relatively sharp peaks at multiples of 60° with additional small peaks in between (Fig. 4A, right). This pattern is clearly distinct from the more sinusoidal modulation of sum grid-cell activity resulting from the conjunctive grid by head-direction cell hypothesis (Fig. 2) and the more full-wave rectified sinusoidal modulation resulting from the repetition suppression hypothesis (Fig. 3).

**Figure 4:**
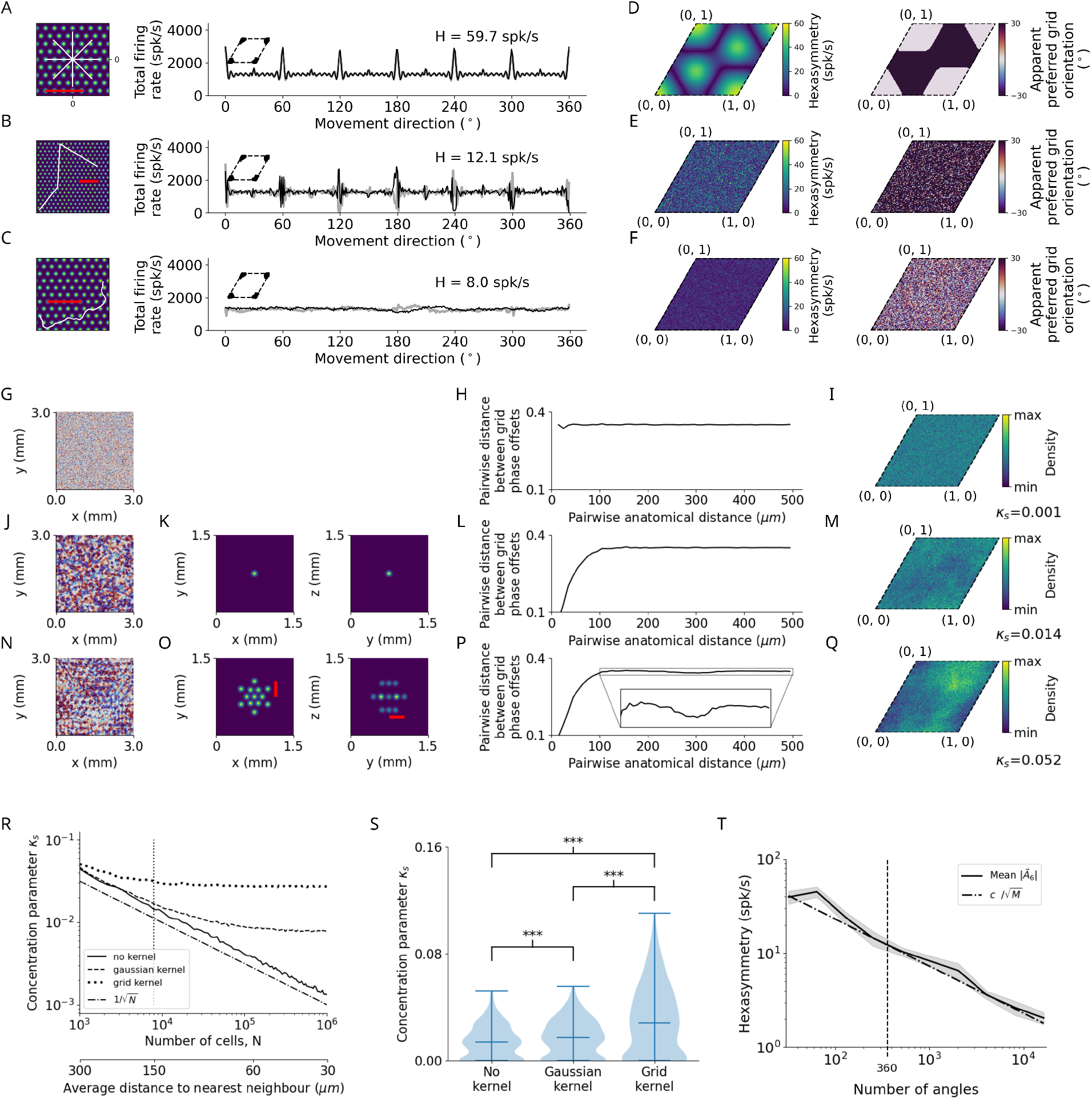
Structure-function mapping hypothesis. (**A–C**) Left: Short example trajectories (white) overlaid onto the firing-rate pattern of an example grid cell (colored). Shown trajectories are for illustration purposes only, and do not reflect the full length of the simulation. The scale bars (red) represent a distance of 120 cm. Right: Population firing rates of 1024 grid cells as a function of the movement direction. Black and light gray lines represent the results from the numerical simulations of Eq. (8) and the analytical derivation in Eq. (32), respectively. Grid phase offsets cells are strongly clustered at (0, 0) with *κ*_*s*_ = 10 (left insets). (**A**) Left: Subsegment of a star-like walk. Right: Population firing rate as a function of the movement direction for star-like runs originating at phase offset (0, 0) with mean firing rate *Ã*_0_ = 1362.4 spk/s (spikes/second) and path hexasymmetry 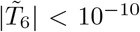. (**B**) Left: Subsegment of a piecewise linear walk. Right: Population firing rate as a function of the movement direction for a piecewise linear trajectory. 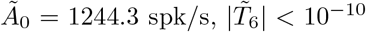. (**C**) Left: Subsegment of a random walk. Right: Population firing rate as a function of movement direction for a random walk. 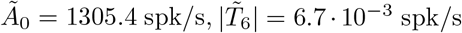 (**D**) Left: Hexasymmetry as a function of the subject’s starting location (relative to grid phase offset). Right: Movement direction associated with the highest sum grid-cell activity, i.e. the phase of peaks in (A, right), has a bimodal distribution (0 or 30°). (**E**) Same as (D) but for piecewise linear walks. (**F**) Same as (D) but for random walks. (**G**) Example of a two-dimensional slice of a three-dimensional (3D) random-field simulation with a spatial resolution of 15 *µ*m. The simulated volume of (3× 3× 3) mm^3^ represents the approximate spatial extent of a voxel in fMRI experiments (for details, see Methods). (**H**) The pairwise phase distance in the rhombus is shown as a function of the pairwise anatomical distance for all pairs of simulated cells from (G). Since no spatial correlation structure is induced, the pairwise phase distance between grid cells remains constant when varying the pairwise anatomical distance between them. (**I**) The resulting phase clustering for 200^3^ simulated grid cells in a (3× 3 ×3) mm^3^ voxel. Brighter colors indicate a higher prevalence of a particular grid phase offset. The distribution of grid phases appears to be homogeneous with a clustering concentration parameter of *κ*_*s*_ = 0.001. (**J–M**) Same as (G–I) but for a convolution of the 3D random field with the 3D Gaussian correlation kernel of width 30 *µ*m shown in (K). Grid cells located next to each other in anatomical space (≲ 30 *µ*m) exhibit similar grid phase offsets (L). No clear clustering is visible for a clustering concentration parameter of *κ*_*s*_ = 0.014 (M). (**N–Q**) Same as (J–M) but for the grid-like correlation kernel with projections shown in (O), which adds some longer-range spatial autocorrelation to the grid phase offsets of different grid cells. The red bars in (O) measures a distance of 300 *µ*m, which corresponds to the separation between two peaks of the correlation kernel lying on the same plane. The pairwise phase distance (P) exhibits a dip around 300 *µ*m. Note in (Q) that the prevalence of particular grid phase offsets is more biased than in (M), with a clustering concentration parameter of *κ*_*s*_ = 0.052. **(R)** Dependence of the clustering concentration parameter *κ*_*s*_ on the number *N* of grid cells in a voxel. A random distribution of grid cells in anatomical space was obtained by subsampling from the 200^3^ grid cells simulated in (G–Q) over 300 realizations. We found *κ*_*s*_ *≈* 0.05 for 10^3^ grid cells in a voxel, and that *κ*_*s*_ decreases monotonically as *N* is increased. Convolution of grid phase offsets with a correlation kernel (as in J–Q) leads to saturation of *κ*_*s*_ for large *N*. Note that the range of values of *κ*_*s*_ here is three orders of magnitude smaller than the strong clustering considered in (A–C). The dashed-dotted line depicts the line 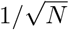 for comparison. The vertical dotted line at *N* = 20^3^ corresponds to the empirically estimated count of grid cells in a (3× 3× 3) mm^3^ fMRI voxel. The secondary lower horizontal axis shows the average distance between regularly distributed grid cells in anatomical space. **(S)** The distribution of the clustering concentration parameter *κ*_*s*_ when using either no kernel, a Gaussian kernel, or a grid-like kernel for 20^3^ grid cells subsampled from the 200^3^ grid cells simulated in (G–Q) over 300 realizations. The grid-like kernel results in a larger maximum value of the clustering concentration parameter, and a large number of realizations results in relatively low clustering. (**T**) The dependence of the hexasymmetry |*Ã*_6_| on the number of angles sampled when unwrapping the star for the piecewise linear walk, averaged over 20 trajectories for each data point. Each additional angle sampled adds 300 cm to the total length of the path. A line proportional to 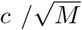 is plotted for comparison, where *c* is an offset parameter (*c* = 4000 spk/s) chosen such that the slope of |*Ã*_6_| can be compared to the slope of 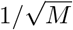, which is proportional to the path hexasymmetry 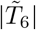 (Eq. 67). The close fit between the solid and dashed lines indicates that the neural hexasymmetry | *Ã*_6_ | is highly correlated with the path hexasymmetry 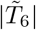 The vertical dotted line at 360 sampled angles corresponds to the number of angles sampled in the star-like run and the piecewise linear run for all main figures.

When examining the structure-function mapping hypothesis for piecewise linear walks and random walks, hexasymmetry appeared to be considerably lower as compared to simulations with star-like walks (Fig. 4, B and C). For piecewise linear walks (but not for random walks), population grid-cell activity as a function of movement direction again exhibited sharp, non-sinusoidal peaks at multiples of 60°, in both positive and negative directions (Fig. 4B). For both piecewise linear walks and random walks, the hexasymmetry values did not show a systematic dependency on the starting location of the subject’s navigation trajectory relative to the locations of the grid fields (Fig. 4, E and F, left), which is due to the fact that these navigation trajectories randomize the starting locations of all path segments. Accordingly, the apparent preferred grid orientations varied randomly as a function of the starting location of the very first path segment and did not exhibit systematic shifts (Fig. 4, E and F, right), which is in contrast to the clear shifts in the apparent preferred grid orientations for star-like walks (Fig. 4D).

In the above-mentioned simulations for the structure-function mapping hypothesis, we chose high values for the clustering of the grid phase offsets from all grid cells. Specifically, we set the clustering parameter to *κ*_*s*_ = 10, due to which the centers of the firing fields of different grid cells were very close to each other (insets in Fig. 4, A–C, right). However, previous empirical studies (Gu et al., 2018; Heys et al., 2014) showed that the clustering parameters (*κ*_*s*_ *≈* 0.1) are orders of magnitude smaller than the strong clustering we considered for our earlier simulations. These empirically determined parameters are actually closer to randomly distributed grid phases as compared to strong, ideal clustering (Fig. 4, G–Q). We thus re-performed our simulations on the structure-function mapping hypothesis using more realistic values, guided by the empirical studies from (Gu et al., 2018; Heys et al., 2014). Specifically, we implemented “simple” clustering using a Gaussian kernel that led to a clustering parameter *κ*_*s*_ *≈* 0.014 (Fig. 4M). Clustering parameters were also low when we added “higher-order” spatial clustering of grid phase offsets by adding some longer-range spatial autocorrelations to the grid-phase offsets (Fig. 4, N– Q) (Gu et al., 2018), which led to a clustering parameter *κ*_*s*_ *≈* 0.052 (Fig. 4Q). These more “realistic” clustering parameters resulted in clearly reduced hexasymmetries.

Specifically, the hexasymmetry was largest (*H* = 59.7 spk/s) for star-like walks with specific starting locations [e.g., (0, 0)], smaller (*H* = 12.1 spk/s) for piecewise linear walks, and smallest (*H* = 8.0 spk/s) for random walks. The type of navigation path thus had a strong influence on the measured neuronal hexasymmetry *H* (Fig. 4A–C). Moreover, the absolute values of these hexasymmetries were quite low compared to the average firing rate of about 1024 spk/s of the simulated population of grid cells (*N* = 1024, each having an average firing rate of 1 spk/s). We thus wondered whether the hexasymmetries of the associated navigation paths substantially contributed to the neuronal hexasymmetries *H*.

In line with this idea, we found that the path hexasymmetries for random walks was proportional to 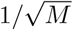 for a large number *M* of steps (Figure S1). We thus examined neural hexasymmetries *H* across a broad range of total trajectory distances (using piecewise linear walks) and observed that *H* decreased also as 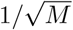 (Fig. 4T). This indicates that the apparent neural hexasymmetry *H* of summed grid-cell activity for piecewise linear walks was driven by random subsamples of all path segments—specifically those path segments crossing through grid fields. These subsamples of path segments necessarily exhibit higher path hexasymmetries than the full set of path segments that has basically zero path hexasymmetry by construction (Fig. S4). We thus conclude that, for piecewise linear walks, hexasymmetry values were driven by a subsampling of the movement directions due to the sparsity of the grid-field locations. Similarly, for a random walk with tortuosity *σ*_*θ*_ = 0.5 rad/s^1/2^, we derived from Figure S1 that the expected path hexasymmetry for 9000 s simulation time (*M* = 0.9 · 10^6^ steps for Δ*t* = 0.01 s) is about 0.007, which results in a contribution to the neural hexasymmetry of *H ≈* 7 spk/s for a population rate of about 1024 spk/s. This number is similar in magnitude to the obtained neural hexasymmetry (*H* = 8.0 spk/s) in Fig. 4C and in Fig. 5 (see also Fig. S4). We thus believe that also for random walks the hexasymmetry *H* obtained in the structure-function mapping case is mainly determined by the path hexasymmetry.

**Figure 5:**
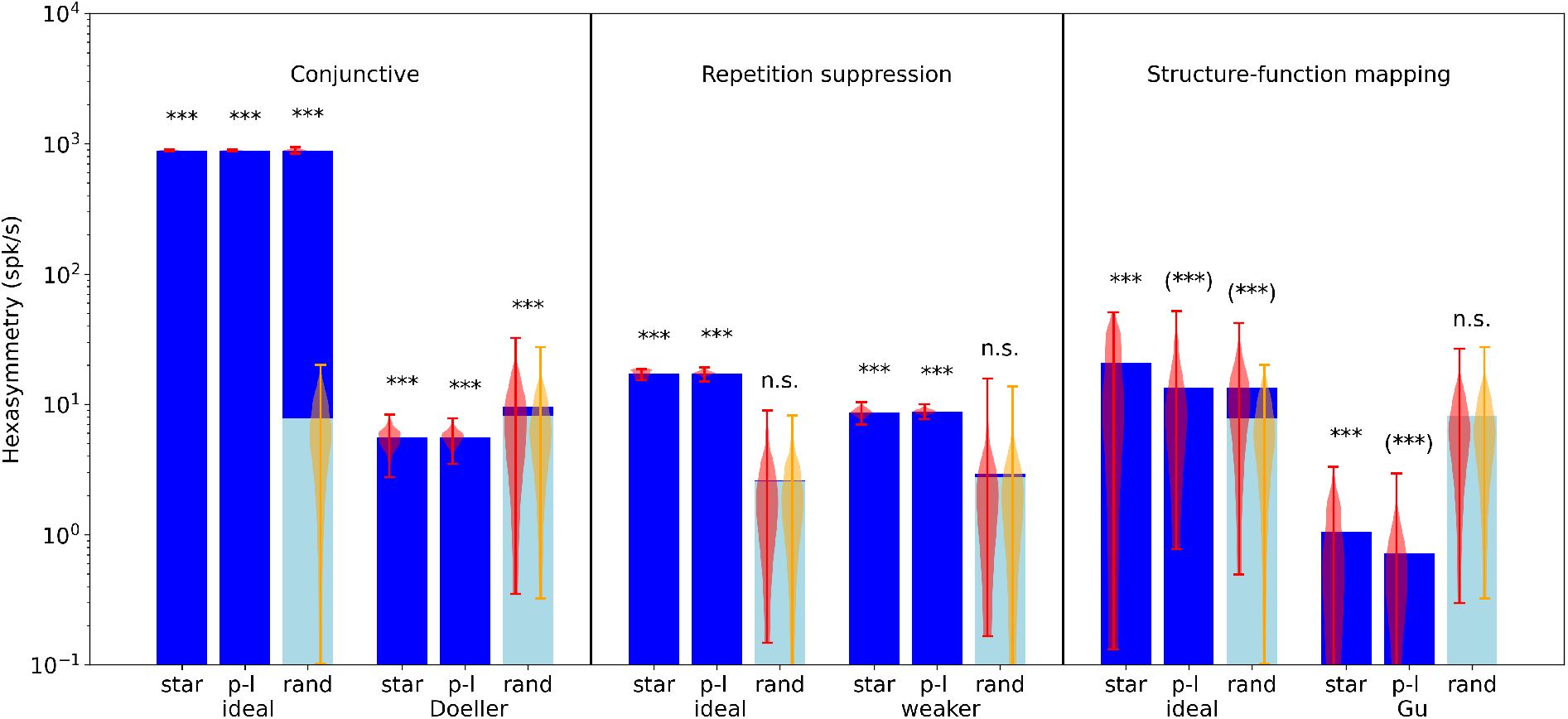
Comparison of hexasymmetry resulting from the three hypotheses. Each of the three hypotheses is implemented for three different types of navigation trajectories: star-like walks (“star”), piecewise linear random walks (“p-l”), and random walks with small step size (“rand”). For each setting, we show the average neural hexasymmetry | *Ã*_6_ | for 1024 cells (dark blue bars) and the average contribution of the trajectory 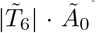 (light blue bars) where *Ã*_0_ is the (variable) average population activity and 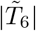 is the hexasymmetry of the trajectory; see Methods for definitions of symbols. The violin plots depict the distributions for firing-rate hexasymmetry (red) and path hexasymmetry (orange). For each hypothesis, we calculate the hexasymmetry for “ideal” parameters (conjunctive: *p*_*c*_ = 1, *κ*_*c*_ = 50 rad^*−*2^, *σ*_*c*_ = 0; repetition suppression: *τ*_*r*_ = 3 s, *w*_*r*_ = 1; clustering: *κ*_*s*_ = 10) as well as more realistic parameters [conjunctive “Doeller”: *p*_*c*_ = 0.33, *κ*_*c*_ = 4 rad^*−*2^, *σ*_*c*_ = 3° motivated by (Doeller et al., 2010; Boccara et al., 2010; Sargolini et al., 2006); repetition suppression “weaker”: *τ*_*r*_ = 1.5 s, *w*_*r*_ = 0.5; clustering “Gu”: *κ*_*s*_ = 0.1 motivated by (Gu et al., 2018)]. For clustering, we also simulated grid phase offsets randomly sampled from a uniform distribution with *κ*_*s*_ = 0 and found that neural hexasymmetries were reduced (compared to “Gu”) by *∼*30% (Fig. S11). The parameters for the random-walk scenario are *T* = 9000 s and Δ*t* = 0.01 s; see Table 1 for a summary of parameters. Each hypothesis condition was simulated for 300 realizations. ***, *P <* 0.001; n.s., not significant; (***), a seemingly significant result (*P <* 0.001) that we think is spurious (see Results section). For pair-wise comparisons of the hexasymmetry values from different trajectory types for each set of parameters, see Fig. S3. For a comparison between our method and previously used methods for evaluating hexasymmetry, see Fig. S8.

Taken together, the structure-function mapping hypothesis with very strong (“ideal”) clustering produces neural hexasymmetry values that are larger than expected from path hexasymmetries only with respect to star-like walks, including a strong dependence on the subject’s starting location (Fig. 4D, left). This range of hexasymmetry values is comparable to those of the repetition-suppression hypothesis with “ideal” parameters (Fig. 3), but values are at least an order of magnitude smaller than in the “ideal” conjunctive grid by head-direction cell case (Fig. 2). When using realistic clustering parameters, neural hexasymmetry values are further reduced as compared to simulations with ideal clustering.

### 2.3 Overall evaluation of the three hypotheses

To provide a systematic evaluation of the three hypotheses, we computed 300 realizations of each hypothesis (using the simulated activity of 1024 cells), separately for each type of navigation and for both ideal and more realistic parameter settings (see Table 1 for a summary of parameter values and section 5.6 “Parameter estimation” for their justification). This resulted in 18 different hypothesis conditions (Fig. 5). For each hypothesis condition, we assessed its statistical significance by performing nonparametric Mann-Whitney U tests between the neural hexasymmetries (*H* := |*Ã*_6_|; see also Eq. 12) and the product 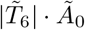 with the multipliers path hexasymmetry 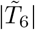 and average population activity *Ã*_0_. For the conjunctive hypothesis, we found that all three types of navigation led to significant neural hexasymmetries. This was the case for both ideal parameters (Mann-Whitney U tests, all *U* = 0, all *P <* 0.001) and for more realistic parameters (Mann-Whitney U tests, all *U* = 0, all *P <* 0.001). For starlike walks and piecewise linear walks, the path hexasymmetries multiplied with the average population activities were near zero because we designed these navigation paths to equally cover all different movement directions. For random walks, the path hexasymmetries multiplied with the average population activities were at values of about 7 spikes per second and showed larger variability because the directions of the random walks were not predefined and could thus vary with regard to their hexasymmetries. If the conjunctive hypothesis was true, fMRI studies should thus see a hexadirectional modulation of entorhinal fMRI activity for all three types of the subjects’ navigation paths.

For the repetition suppression hypothesis, we found that star-like walks and piecewise linear walks resulted in significant neural hexasymmetries for both ideal and weaker parameters (Mann-Whitney U tests, all *U* = 0, all *P <* 0.001), whereas random walks did not (Mann-Whitney U tests, both *U >* 1753, both *P >* 0.405). For random walks, the path hexasymmetries multiplied with the average population activities were lower as compared to the other two hypotheses because the repetition suppression necessarily leads to lower average population activities. If the repetition suppression hypothesis was true, fMRI studies should thus observe significant neural hexasymmetries only for star-like walks and piecewise linear walks with long enough segments, whereas random walks (with large enough tortuosities, Fig. S2) should not lead to significant neural hexasymmetries.

Regarding the structure-function mapping hypothesis, the statistical tests showed that most of the hypothesis conditions resulted in significant neural hexasymmetries as compared to the path hexasymmetries multiplied with the average population activities. This was the case for both ideal parameters

(Mann-Whitney U tests, all *U <* 736, all *P <* 0.001) and more realistic parameters (Mann-Whitney U tests for star-like walks and piecewise linear walks, all *U* = 0, all *P <* 0.001). Only the hypothesis condition with random walks and more realistic parameters exhibited no significant neural hexasymmetries (Mann-Whitney U test, *U* = 1786, *P* = 0.472). However, these results regarding the structure-function mapping hypothesis should be treated with great caution. Firstly, the neural hexasymmetries for starlike walks heavily depend on the starting location of the subject relative to the grid fields, and different starting locations lead to different apparent grid orientations (Fig. 4D). Secondly, the significant results for the navigation conditions with piecewise linear walks and random walks actually result from an inhomogeneous sampling of movement directions through the grid fields and therefore do not reflect true neural hexasymmetries (Fig. 4T). Only the structure-function mapping hypothesis is susceptible to this effect because the grid fields are clustered at similar spatial locations (whereas the grid fields are homogeneously distributed in the case of the conjunctive hypothesis and the repetition suppression hypothesis). In simulations with infinitely long paths, the neural hexasymmetries (for the navigation types of piece-wise linear walks and random walks) would not be significantly higher than the path hexasymmetries multiplied with the average population activities. In empirical studies, this effect can be detected by correlating the subject-specific path distances with the subject-specific neural hexasymmetries: if there is a generally negative relationship, this will hint at the fact that the neural hexasymmetries are basically due to relevant path hexasymmetries of path segments crossing the grid fields. In essence, we therefore believe that the structure-function mapping hypothesis leads to true neural hexasymmetry only in the case of star-like walks.

### 2.4 Influence of other factors

Our simulations above were performed in an infinite spatial environment, which is different from empirical studies in which subjects navigate finite environments. We were thus curious whether the size and shape of finite environments could affect the strength of hexadirectional modulations of population grid-cell activity.

Our simulations showed that for both circular and square environments, hexasymmetry strengths did not considerably depend on the size of the environment when the subject performed random walks (Fig. S5). Similarly, rotating the navigation trajectories relative to the grid patterns did not affect the hexasymmetry strengths (Fig. S6). These results thus suggest that experiments in animals and humans can use various types and sizes of the environments to investigate hexadirectional modulations of sum grid-cell activity.

## 3 Discussion

We performed numerical simulations and analytical estimations to examine how the activity of grid cells could potentially lead to a neural population signal in the entorhinal cortex. Such a neural population signal has been observed in multiple fMRI studies including (Doeller et al., 2010; Kunz et al., 2015; Constantinescu et al., 2016; Horner et al., 2016; Bellmund et al., 2016; Stangl et al., 2018; Nau et al., 2018; Julian et al., 2018; Bierbrauer et al., 2020; Julian and Doeller, 2021; Bongioanni et al., 2021; Moon et al., 2022) and iEEG/MEG studies including (Maidenbaum et al., 2018; Chen et al., 2018; Staudigl et al., 2018; Chen et al., 2021; Wang and Wang, 2021; Convertino et al., 2023), and consists of a hexadirectional modulation of the entorhinal fMRI/iEEG signal as a function of the subject’s movement direction through its spatial environment. We note though that some studies did not find evidence for neural hexasymmetry. For example, a surface EEG study with participants “navigating” through an abstract vowel space did not observe hexasymmetry in the EEG signal as a function of the participants’ movement direction through vowel space (Kaya et al., 2020). Another fMRI study did not find evidence for grid-like representations in the ventromedial prefrontal cortex while participants performed value-based decision making (Lee et al., 2021). This raises the question whether the detection of macroscopic grid-like representations is limited to some recording techniques (e.g., fMRI and iEEG but not surface EEG) and to what extent they are present in different tasks.

### 3.1 Possibility of other mechanisms underlying hexadirectional population signals in the entorhinal cortex

We examined three hypotheses that have been previously suggested as potential mechanisms underlying the emergence of the hexadirectional population signal in the entorhinal cortex (Doeller et al., 2010; Kunz et al., 2019) and found that all three hypotheses can—in principle and in ideal situations—lead to a hexadirectional modulation of entorhinal cortex population activity. However, none of the three hypotheses described here may be true and that another mechanism may explain macroscopic grid-like representations. This includes the possibility that neural hexasymmetry is completely unrelated to grid-cell activity, previously summarized as the ‘independence hypothesis’ (Kunz et al., 2019). For example, a population of head-direction cells whose preferred head directions occur at offsets of 60 degrees from each other could result in neural hexasymmetry in the absence of grid cells (though such a case would require an explanation for why the preferred head directions of different cells are offset by 60 degrees).

### 3.2 Navigation paths have a major influence on the hexadirectional population signal

A major observation of this study is that the way how subjects navigate through the environment has a major influence on whether a hexadirectional population signal can be observed. We distinguished three major types of navigation: navigation with random walk trajectories in which straight paths are quite short, which resembles the navigation pattern in rodents; navigation with piecewise linear trajectories in which the subject navigates along straight paths combined with random sharp turns between the straight segments; and navigation with star-like trajectories in which the subject starts each path from a fixed center location and navigates along a straight path with a predetermined allocentric direction and distance. Critically, we found that the conjunctive hypothesis leads to a hexadirectional population signal irrespective of the specific type of navigation; that the repetition suppression hypothesis leads to hexadirectional population signals only in the case of star-like trajectories and piecewise linear trajectories (but not for random trajectories); and that for the structure-function mapping hypothesis “true” hexadirectional population signals can only be observed for star-like trajectories.

The finding that the type of navigation paths influences whether a hypothesis can lead to hexadirectional population signals of the entorhinal cortex is informative to future fMRI/iEEG studies, which could empirically evaluate which of the three hypotheses is most likely to be true: By asking or requiring the subjects to navigate in different ways through the task environments (in a star-like fashion, in a piecewise linear fashion, and in a random fashion), these future fMRI/iEEG studies could demonstrate whether hexadirectional population signals are present in all three navigation conditions (speaking in favor of the conjunctive hypothesis); whether they are present only during star-like or piecewise linear trajectories (in favor of the repetition suppression hypothesis); or whether they are mainly visible for star-like trajectories and exhibit a systematic decrease with increasing total trajectory length for piecewise linear and random walks (in this case speaking in favor of the structure-function mapping hypothesis). We note that in our simulations, each linear path segment of a piecewise linear walk has a length that is ten times larger than the grid scale. While humans performing navigation tasks exhibit straighter trajectories than those of rats (Doeller et al., 2010; Kunz et al., 2015; Horner et al., 2016), they are still more similar in scale to random walks than piecewise linear walks. Thus neural hexasymmetries were only significant for the conjunctive hypothesis when using empirical navigation paths from previous studies (Fig. S7). In contrast to this major effect of navigation type on the presence of hexadirectional signals, the size and shape of the environment did not influence the strength of the hexadirectional signals in a relevant manner (Figs. S5 and S6).

The structure-function mapping hypothesis predicts that in fMRI studies involving star-like walks, there may be up to 30° shifts in the orientation of hexadirectional modulation between neighboring voxels (Fig. 4D). This is in contrast with the other two hypotheses, where the preferred grid orientations are similiar between voxels, and may differ only slightly when recording distinct grid modules with different grid orientations (Stensola et al., 2012).

### 3.3 A note on our choice of the values of model parameters

Another insight of this study is that the exact biological properties of grid cells play a major role regarding the question whether hexadirectional population signals can be observed. For example, the conjunctive hypothesis cannot lead to hexadirectional population signals if the tuning width of the conjunctive grid by head-direction cells is too broad (Fig. 2E). Here, the percentage of strongly tuned conjunctive cells also plays a relevant role. Empirical studies in rodents found a range of tuning widths among grid cells ranging from broad to narrow Doeller et al. (2010); Sargolini et al. (2006). The percentage of conjunctive cells in the entorhinal cortex with a sufficiently narrow tuning may thus be low. Such distributions (with a proportionally small amount of narrowly tuned conjunctive cells) lead to a systematic decrease in the absolute hexasymmetry values. The neural hexasymmetry in this case would be driven by the subset of cells with sufficiently narrow tuning widths. If this causes the neural hexasymmetry to drop below noise levels, the statistical evaluation of this hypothesis would change.

We also found that hexadirectional population signals only emerge if the preferred directions of the conjunctive cells are aligned precisely enough with the grid axes of the grid cells (Fig. 2E). Furthermore, no hexadirectional population activity will emerge if the preferred head directions of conjunctive grid-by-head-direction cells exhibit other types of rotational symmetry such as 10-fold rotational symmetry reported in recordings from rats (Keinath, 2016). Additionally, while we assumed that all conjunctive grid cells maintain the same preferred head direction between different firing fields, conjunctive grid cells have also been shown to exhibit different preferred head directions in different firing fields (Gerlei et al., 2020). This could lead to hexadirectional modulation if the different preferred head directions are offset by 60° from each other, but will not give rise to hexadirectional modulation if the preferred head directions are randomly distributed. To the best of our knowledge, the distribution of preferred head directions was not quantified by Gerlei et al. (2020), thus this remains an open question. Together, it currently seems possible that grid cells and conjunctive grid by head-direction cells meet the necessary biological properties for the conjunctive hypothesis (Doeller et al., 2010), but further studies on the characteristics of conjunctive grid by head-direction cells (also in humans) will be useful.

The exact biological properties of grid cells play also a major role for the structure-function mapping hypothesis, which heavily relies on the property of neighboring grid cells to share a similar grid phase offset (i.e., a high spatial autocorrelation of grid phases) and whether there might be longer-range spatial autocorrelations between the grid phases (Gu et al., 2018). With the evidence at hand, it seems unlikely to us that the grid phases are clustered strongly enough to facilitate a hexadirectional population signal. Specifically, we found that the clustering parameters of grid phase offsets gleaned from existing empirical studies (Gu et al., 2018) produced the smallest neural hexasymmetry when compared to the other conditions we considered, while the neural hexasymmetries obtained from simulations randomly sampling grid phase offsets from a uniform distribution on the unit rhombus (that one might refer to as the ‘standard grid cell model’) were only slightly smaller.

Regarding realistic parameters for the repetition suppression hypothesis, we are currently not aware of detailed measurements of the relevant grid-cell properties (i.e., the adaptation time constant *τ*_*r*_ and the adaptation weight *w*_*r*_) so that it remains unclear to us to what extent the repetition suppression hypothesis is biologically plausible. Future studies may thus quantify the relevant properties of (human) grid cells in greater depth to help clarify which hypothesis regarding the emergence of hexadirectional population signals may most likely be true.

### 3.4 Influence of grid scale on hexasymmetry

In this study, we assumed that all grid cells have the same grid spacing of 30 cm. For rats, this is at the lower end of grid spacing values, corresponding to the dorsal region of the medial EC (Stensola et al., 2012). How would the three hypotheses perform for larger values of grid spacing? The conjunctive hypothesis would remain unchanged as it does not depend on grid spacing. In contrast, the repetition suppression hypothesis depends on the grid scale. Here, the strongest hexasymmetry is achieved when the adaptation time constant is similar to the time that it takes the subject to move between neighboring grid fields (Fig. 3B). Altering the grid scale changes this traversal time between grid fields and thus hexasymmetry. In Fig. 3, we chose the “ideal” adaptation time constant to maximize hexasymmetry. A change in grid scale would thus reduce hexasymmetry for this particular value of the adaptation time constant. As there are different grid-cell modules in the entorhinal cortex that exhibit different grid spacings (Stensola et al., 2012), repetition-suppression effects may vary across modules (if the adaptation time constant is not tuned in accordance with the grid scale). Regarding the structure-function mapping hypothesis, different grid spacings can have a major effect on neural hexasymmetry if the path segments are short compared to the grid scale, similar to effects of the starting location (Fig. 4D). For longer path segments, altering the grid scale should have a smaller effect on neural hexasymmetry. Random walks and piecewise-linear walks do not lead to real neural hexasymmetries, and different grid spacings are thus irrelevant. Overall, different grid spacings are thus most important in the context of the repetition-suppression hypothesis.

Another factor that might complicate the picture is an interaction of different grid scales: rodent grid cells have been found to be organized into 4–5 discrete modules with different grid scales (Stensola et al., 2012) and this may also be true in humans. Together with the anatomical dorsoventral length of the human entorhinal cortex (*≈* 2 cm, Behuet et al. (Springer International Publishing, 2021)), this means that a module might cover roughly 400–500 *µ*m. A typical voxel of 3 × 3 × 3 mm^3^ would thus either comprise one grid module or more than one module. The case of a voxel comprising a single module was already discussed. If a voxel comprised two or more modules, there would be at least two subpopulations of grid cells with different grid spacings.

For the structure-function mapping hypothesis, significant neural hexasymmetry can be achieved only for star-like trajectories. In this scenario, the hexasymmetry depends on the starting position of the trajectories, i.e. the center position of the star (Fig. 4D). Two subpopulations of grid cells from different modules essentially act like two individual grid cells in the context of the structure-function mapping hypothesis. Thus, they can either add their hexasymmetry values (e.g. when the star center is in the middle of a grid field for both subpopulations) or cancel out (when the star center is in a regime of 0° grid phase for one subpopulation and in a regime of *±* 30° grid phase for the other).

For the repetition suppression hypothesis, two populations of grid cells with the same adaptation time constant but different grid spacing would result in two different values of hexasymmetry, depending on the interaction between grid-field traversal time and adaptation time constant. The joint hexasymmetry value would then (i) be smaller than if all grid cells had the same grid spacing (matched to the adaptation time constant) and (ii) depend on the population sizes. We note that the adaptation time constant might be tuned to match the grid spacing. This would result in high hexasymmetry values for both subpopulations, and thus also for the full population.

### 3.5 Sources of noise

The neural hexasymmetry as defined by Eq. (12) also contains contributions from the path hexasymmetry of the underlying trajectory. Star-like walks and piecewise linear random walks have a path hexasymmetry of zero by construction. Therefore, these trajectory types do not contribute to the neural hexasymmetry. For random walks, the path hexasymmetry depends on the number of time steps *M* and the tortuosity parameter *σ*_*θ*_ (Fig. S1). To compare the path hexasymmetry to the neural hexasymmetry, the path hexasymmetry needs to be multiplied by the time-averaged population activity. Since the path hexasymmetry ranges from zero to one (Eq. 14), its contribution to the neural hexasymmetry ranges from zero (for a uniform sampling of movement directions) to the time-averaged population activity (if there is only one movement direction). For *M* = 900 000 time steps and a tortuosity of *σ*_*θ*_ = 0.5 rad/s^1/2^, random walk trajectories contribute *∼* 0.8% of the time-averaged population activity to the neural hexasymmetry (Fig. S1).

Another factor that contributes noise is the finite sampling of grid phase offsets. For all our simulations of the conjunctive grid by head-direction cell hypothesis and the repetition suppression hypothesis, we sample a finite number of grid phase offsets from a two-dimensional von Mises distribution with a clustering parameter *κ*_*s*_ = 0 (equivalent to a uniform distribution on the unit rhombus). In this scenario, random fluctuations give rise to a non-zero clustering parameter *κ*_*s*_ with a corresponding mean neural hexasymmetry of 0.7 spk/s (labeled “standard” and “star” in Fig. S11). For 1024 grid cells, this contribution is ten times smaller than the neural hexasymmetry resulting from the ‘‘realistic” parameters for both the conjunctive grid by head-direction cell hypothesis and the repetition suppression hypothesis (Fig. 5). In the case of the structure-function mapping hypothesis, these fluctuations would not be considered as ‘noise’ as they directly contribute to the hexasymmetry from the clustering of grid phase offsets.

### 3.6 On the relation of firing rates and fMRI/iEEG signals

A topic that this study did not investigate is the question of how the sum signal of single neurons translates into fMRI and iEEG signals. In neocortical regions such as the auditory cortex, a clear linear relationship between single-neuron activity and fMRI activity has been observed (Mukamel et al., 2005), but it remains elusive whether this linear relationship also applies to the entorhinal cortex in general and to entorhinal grid cells in particular. In the neighboring hippocampus, for example, the relationship between single-neuron activity and the fMRI signal is highly complex (Ekstrom, 2010; Kunz et al., 2019). Future studies are needed to detail the relationship between single-neuron firing and fMRI/iEEG signals in the entorhinal cortex. This would allow us to clarify whether a hexadirectional modulation of sum grid-cell activity directly results in a hexadirectional modulation of fMRI/iEEG activity or whether currently unknown factors modulate the expression of hexadirectional fMRI/iEEG signals.

## 4 Conclusion

Using numerical simulations and analytical derivations we showed that a hexadirectional neural population signal can emerge from the activity of grid cells given the ideal conditions of three different hypotheses. Whether a given hypothesis leads to a hexadirectional population signal is significantly influenced by the subjects’ type of navigation through the spatial environment and by the exact biological properties of human grid cells.

## 5 Methods

### 5.1 Trajectory modeling

To describe grid-cell activity as a function of time *t* during which a subject (animal or human) is exploring an environment, we model three distinct trajectory types. We first describe trajectory types in environments without bounds, which are quasi infinite, and then add rules that account for boundaries.

#### 5.1.1 Environments without boundaries

The first trajectory type is a random walk *X*_*t*_ = [*x*_*t*_, *y*_*t*_], which is defined by

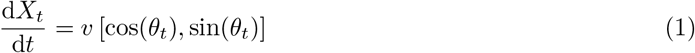

with *θ*_*t*_ = *σ*_*θ*_ *· W*_*t*_ where *σ*_*θ*_ controls the tortuosity of the trajectory and *W*_*t*_ is a standard Wiener process. In a numerical simulation with a time step Δ*t*, the angle is updated in each time step by *θ*_*t*+Δ*t*_ = *θ*_*t*_ + *σ*_*θ*_ · Δ*W* where Δ*W* is a normally distributed random variable with variance Δ*t*. The variable *v* depicts the (constant) speed.

The second type of navigation is a star-like walk, where the subject moves radially outwards from a predefined origin in space at a certain angle *θ* on a straight line to a maximum distance *r*_*max*_ at a constant speed *v*. In simulations, this movement is repeated (with the same predefined origin) for *N*_*θ*_ angles that are equally spaced on the interval [0, 2*π*). Within each individual radial path, the subject does not turn around and move back to the origin, i.e., the entire trajectory of *N*_*θ*_ radial paths is not continuous.

Finally, we introduce a piecewise linear walk, which is constructed by placing all the radial paths of the star-like walk end-to-end such that they form one single continuous trajectory of length *N*_*θ*_ *r*_*max*_. The trajectory thus consists of successive straight runs for the simulated subject, which can be interpreted as a random walk with a time step Δ*t* = *r*_*max*_*/v* and directions that are sampled uniformly without replacement from a predetermined set of angles. In comparison to the random walk and the star-like walk, this procedure presumably reflects the situation in human virtual-reality setups most closely, as participants often move along straight trajectories with intermittent turns (Doeller et al., 2010; Kunz et al., 2015; Horner et al., 2016).

#### 5.1.2 Environments with boundaries

Most virtual-reality studies in humans use finite instead of infinite spatial environments to examine grid-like representations. We wondered whether the size and the shape of these finite environments might modulate the strength of macroscopic grid-like representations obtained through one or more of the three hypotheses. Hence, we performed our simulations not only in infinite but also in finite environments with a given size (between one and six times the grid spacing) and shape (circle and square).

For random-walk trajectories, we enforce that the navigating subject stays within the circular or square environment by performing an “out-of-bounds check” at each time point. This means that, after every time step Δ*t*, we measure the distance that the subject has moved outside of the boundary. This is done differently in square and circular environments, both of which are centered at the origin (*x* = 0, *y* = 0). In the square environment, we define the variables Δ*x* and Δ*y* as the distance the subject has moved out of bounds in the *x* and *y* coordinates, respectively. Let *L* be the half of the length of its sides. Δ*x* and Δ*y* are then defined as Δ*x* = max [|*x*| *−L*, 0] and Δ*y* = max [|*y*| *−L*, 0]. For circular environments, let *R* be the radius of the circle, and let 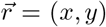 be the position vector of the subject. We then introduce the measure 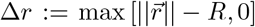, such that Δ*r* is non-zero only when the subject has moved outside of the circular boundary. If at any time point either Δ*x*, Δ*y*, or Δ*r* are non-zero, the out-of-bounds check fails. In this case, we reject the movement in this time step and keep resampling a new angle *θ*_*t*_ (Kropff and Treves, 2008; Si et al., 2012) until the check succeeds, meaning that the subject has made a move that is within the boundaries of the environment. If Δ*x*, Δ*y*, or Δ*r* remain non-zero for 50 consecutive samples of *θ*_*t*_, we temporarily increase the tortuosity *σ*_*θ*_ by a factor 1.1. Without this increase in tortuosity, the subject tends to get stuck when approaching the boundary at angles close to the perpendicular or at the corners of the square boundary. The tortuosity is reset to its initial value once a valid move is made. We visually checked the random-walk trajectories, which show some oversampling along the boundaries, and found that they were comparable to the navigation trajectories in rodent studies (e.g. Hafting et al., 2005).

### 5.2 Implementation of grid cell activity

The activity profile *G*_*i*_ of grid cell *i* (for *i* = 1, …, *N* in a population of *N* grid cells) is modelled as the product of three cosine waves rotated by 60° (= *π/*3) from each other:

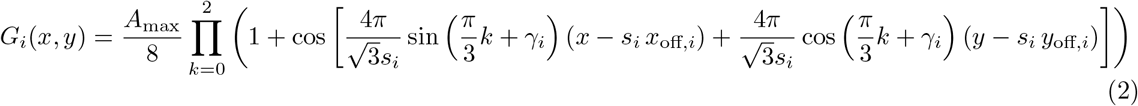

where *A*_max_ is the grid cell’s maximal firing rate, *s*_*i*_ depicts the cell’s grid spatial scale (“grid spacing”), *x*_off,*i*_ and *y*_off,*i*_ are the phase offsets of the grid (“grid phase”) in the two spatial dimensions (called *x* and *y* here), and *γ*_*i*_ is the orientation of the grid (“grid orientation”); see Table 1 for numerical values of parameters and Fig. 1A for an illustration of the three grid characteristics.

To describe the activity of many grid cells (for example in a voxel for an MRI scan), we sum up the firing rates of *N* grid cells with the same grid spacing and orientation but different offsets in Eq. (2). For a given trajectory *X*_*t*_ = [*x*_*t*_, *y*_*t*_], the macroscopic activity as a function of time *t* is then simply described _*i*=1_ by the sum 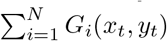.

### 5.3 Implementation of the three hypotheses to explain macroscopic grid-like representations

Here we summarize how the activity in a population of *N* grid cells can be described if they also exhibit (i) head-direction tuning, (ii) repetition suppression (i.e., firing-rate adaptation), or (iii) grid phases that are clustered across grid cells.

In all our models, the activity *G*_*i*_ of a grid cell is described by Eq. (2). Different grid cells typically have different phase offsets (*x*_off,*i*_, *y*_off,*i*_) but the same grid spacing *s*_*i*_ := *s*, ∀*i* and grid orientation *γ*_*i*_ := *γ*, ∀*i* (Hafting et al., 2005; Boccara et al., 2010; Gardner et al., 2022).

#### 5.3.1 Conjunctive grid by head-direction cell hypothesis

To include head-direction tuning in our model, we note that a given trajectory *X*_*t*_ has an angle *θ*_*t*_ at time *t*. The summed firing rate, i.e. the population activity, *A*^*c*^ from *N* such conjunctive cells can then be described by

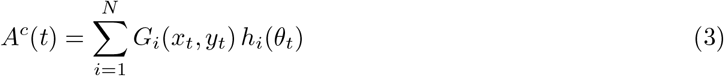

where the upper index ‘c’ indicates “conjunctive” and where we incorporate conjunctive grid-head direction (HD) tuning via the (scaled by factor 2*π*) von-Mises distribution

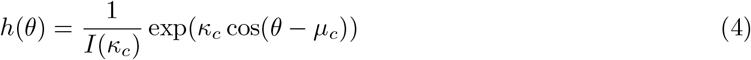

with concentration parameter *κ*_*c*_ and preferred angle *µ*_*c*_. The symbol *I* represents the modified Bessel function of the first kind of order 0. The parameter *κ*_*c*_ describes the width of the HD tuning: if *κ*_*c*_ *→− ∞*, the HD tuning is sharpest; the smaller *κ*_*c*_, the wider the HD tuning (see Fig. 2); for *κ*_*c*_ = 0, there is no HD tuning, and our scaling leads to *h*(*θ*) *≡* 1. We choose the preferred angle as 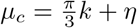 where *k* is randomly drawn from *{*0, 1, 2, 3, 4, 5*}* and *η* is randomly drawn from a normal distribution with mean 0 and standard deviation *σ*_*c*_. For *σ*_*c*_ = 0, the directional tuning is thus centered around a multiple of 60°. The parameter *σ*_*c*_ introduces jitter in the alignment of directional tuning to one of the grid axes.

We modelled the cases in which all grid cells show HD tuning (“ideal” case, fraction of conjunctive cells: *p*_*c*_ = 1) as well as a more “realistic” case in which only a third of the cells is conjunctive (*p*_*c*_ = 1/3; (Boccara et al., 2010; Sargolini et al., 2006)). We note that this is an approximation, since the proportion of conjunctive cells is highly variable across layers of the entorhinal cortex, with up to 90% conjunctive cells in layer V.

#### 5.3.2 Repetition suppression hypothesis

To incorporate repetition suppression in the model, we add an explicit dependence of grid-cell activity on time *t*. Specifically, we subject the firing rate *G*_*i*_ of a grid cell to an adaptation mechanism:

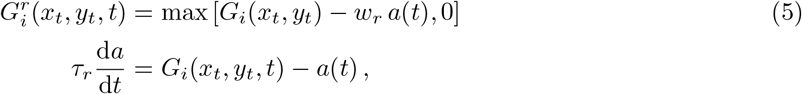

where *a* depicts the adaptation variable, and *τ*_*r*_ and *w*_*r*_ are the repetition-suppression time constant and the weight of the suppression, respectively. The upper index ‘r’ in 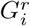 indicates “repetition suppression”. The adaptation time constant *τ*_*r*_ is usually on the order of seconds. We restrict the adaptation weight *w*_*r*_ to the interval [0, 1] as negative values would cause enhancement rather than suppression, and values larger than one would lead to suppression that is stronger than the peak activity of the single grid cell (Fig. 3B). The maximum operation ‘max(*x*)’ in Eq. (5) ensures that the output firing rate 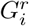 is always positive. Together, the summed firing rate *A*^*r*^ from *N* such adapting cells can then be described by

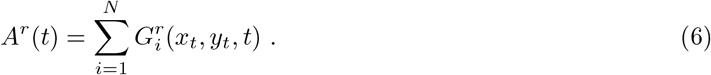

We note that the explicit dependence of the firing rate 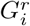 of grid cell *i* on time *t* needs to be considered separately for every cell for repetition suppression, which makes numerical simulations more computationally expensive. In contrast, the functions *G*_*i*_ (in Eq. (3)) and *h*_*i*_ (in Eq. (4)) for the conjunctive hypothesis depend also on time but only implicitly via the location [*x*_*t*_, *y*_*t*_] and the direction *θ*_*t*_ of movement at time *t* — and therefore the explicit time dependence of individual cells can be disregarded, which makes numerical simulations computationally cheaper.

#### 5.3.3 Structure-function mapping hypothesis

The structure-function mapping hypothesis relies on a preferred grid phase for neighboring cells. We use two possible choices for the set of grid phases (*x*_off,*i*_, *y*_off,*i*_): they are either drawn from a uniform distribution on the unit rhombus or clustered. For clustered spatial phases, we draw *x*_off,*i*_ and *y*_off,*i*_ independently, each from a univariate von-Mises distribution (with a defined central phase *µ*_*s*_ and concentration parameter *κ*_*s*_). For a grid phases drawn from a uniform distribution, we note that random fluctuations lead to a certain degree of clustering of the grid-phase sample 11. We can describe the resulting summed activity of *N* grid cells simply by

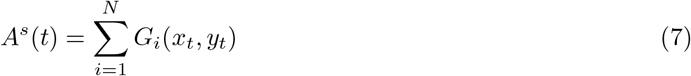

with the upper index ‘s’ representing “structure-function”.

### 5.4 Quantification of hexasymmetry of neural activity and trajectories

Combining the mathematical descriptions of grid cell activity for the three hypotheses (“conjunctive”, “repetition suppression”, and “structure-function”), we can denote the resulting population activity *A* of *N* grid cells by

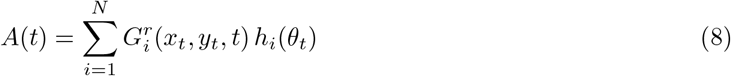

where 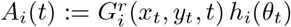 is the firing rate of cell *i*. To derive from *A* the activity as a function of movement (or heading) direction *θ*_*t*_, we focus on time steps of length Δ*t* in which the trajectory is linear. In time step *m*, i.e., for time *t* in the time interval [*t*_*m*_, *t*_*m*+1_) where *t*_*m*_ = *m*Δ*t* and *m* is an integer, the trajectory has the fixed angle *θ*_*m*_. The time-discrete mean activity *Ā*(*t*_*m*_) associated to this interval is the average of *A*(*t*) along the linear segment of the trajectory:

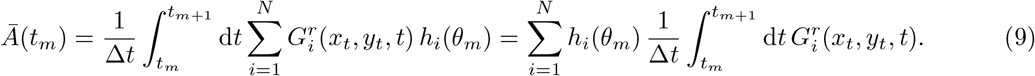

The integral in Eq. (9) is either calculated analytically, as derived in the following section, or numerically. For a total number of *M* time steps in a trajectory, the (normalized) mean activity *Ã*(*ϕ*) as a function of some head direction *ϕ* is then

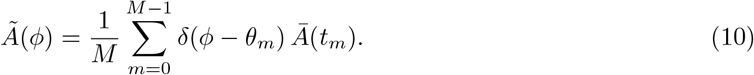

where *δ* is the Dirac delta distribution. With complex Fourier coefficients *c*_*n*_ (with *n ∈* ℕ) defined as

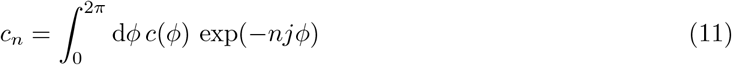

we can quantify the hexasymmetry *H* of the activity of a population of grid cells as

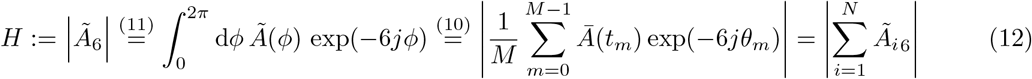

where *Ã*_*i*6_ is the 6th Fourier coefficient of cell *i*. The hexasymmetry is nonnegative (*H >*= 0) and it has the unit spk/s (spikes per second). Furthermore, the average (over time) population activity can be expressed as

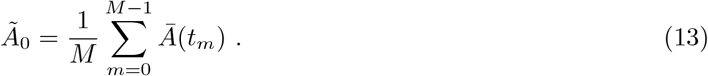

The hexasymmetry *H* could be generated by various properties of the cells, but *H* as defined above contains also contributions from the hexasymmetry of the underlying trajectory. This is due to the fact that we sum up in Eq. (10) the population activities *Ā*(*t*_*m*_) without taking into account the distribution of movement directions *θ*_*m*_. In this way, a hexasymmetry that is potentially contained in the subject’s navigation trajectory contributes to the hexasymmetry *H* of the neural activity.

Empirical (fMRI/iEEG) studies (e.g. Doeller et al., 2010; Kunz et al., 2015; Maidenbaum et al., 2018) addressed this problem of trajectories spuriously contributing to hexasymmetry by fitting a Generalized Linear Model to the time discrete mean activity. In contrast, our new approach to hexasymmetry in Eq. (12) is able to quantify the contribution of the path to the neural hexasymmetry, and has the advantage that it allows an analytical treatment (see next section). Comparing our new method with previous methods for evaluating hexasymmetry led to qualitatively identical statistical effects, but methods which use binning (such as in Kunz et al. 2015) led to decreased effect sizes (Fig. S8).

To nevertheless be able to quantify how much a specific trajectory contributed to the neural hexasymmetry *H* =| *Ã*_6_|, we also explicitly quantified the hexasymmetry of navigation trajectories and interpreted *H* relative to the path hexasymmetry. To quantify the hexasymmetry of a trajectory, we used the same approach as in Eq. (10) and defined the distribution of movement directions of the trajectory by

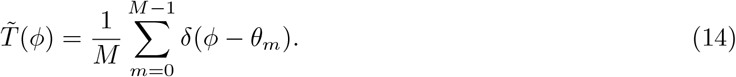

The path hexasymmetry of the trajectory is then 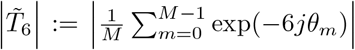, which is similar to Eq. (12). The path hexasymmetry is a unitless quantity, and ranges from zero (for a uniform sampling of movement directions) to one (when only one movement direction is sampled). To be able to estimate how much of the hexasymmetry *H* of neuronal activity is due to the hexasymmetry of the trajectory, we compare the relative hexasymmetry of the activity, |*Ã*_6_ */Ã*_0_|, with the relative hexasymmetry of the trajectory, 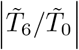, noting that 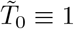. The two terms being similar in magnitude, i.e. 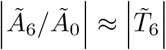, indicates that the trajectory is a major source of hexasymmetry whereas 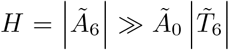 suggests that hexasymmetry has a neural origin.

#### 5.4.1 Analytical derivation of mean activity

In the following, we provide derivations that allow us to analytically integrate Eq. (9) for the conjunctive grid by head-direction cell hypothesis and the structure-function mapping hypothesis but not for the repetition suppression hypothesis. The respective results are shown in Fig. 2B–H and Fig. 4A–C, demonstrating that they are very similar to the numerical results.

We analytically calculate the mean activity *Ā* by averaging *A* along a linear segment of a trajectory (cf. Eq. (9)). For convenience, the following abbreviations are used in Eq. (2) with the same grid spacing, *s*_*i*_ = *s*, and the same grid orientation, *γ*_*i*_ = *γ*, for all cells *i*:

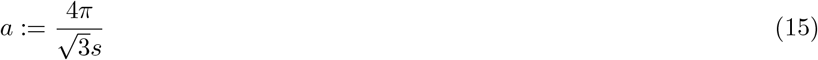

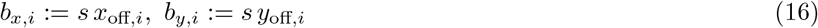

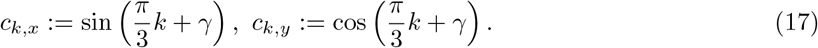

##### A single grid cell

We start with a single grid cell without head-direction tuning and without repetition suppression. Eq. (2) can be described in polar coordinates

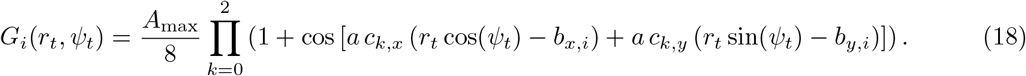

In order to integrate along a piece of a straight line through the origin (similarly to the star-like walk), the angle 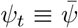 can now be kept fixed (for that particular straight line) and we only have to consider 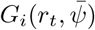. If we define *r*_*m*_ and *r*_*m*+1_ as the distances from zero that the subject is located at at times *t*_*m*_ and *t*_*m*+1_ respectively, integration by substitution gives us

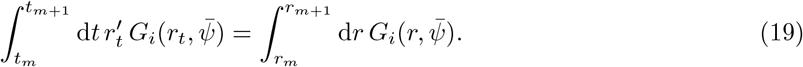

Since the speed of movement 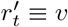 is assumed to be constant along the whole trajectory and Δ*r* = *v*Δ*t*, for Δ*r* := *r*_*m*+1_ *− r*_*m*_, we obtain

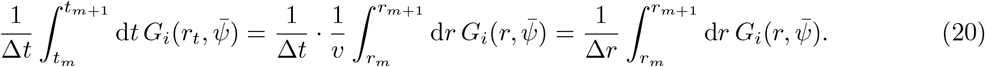

Thus, we have

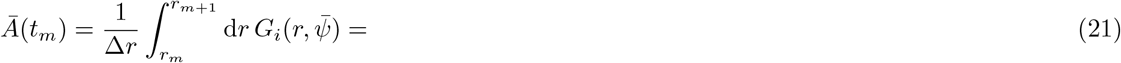

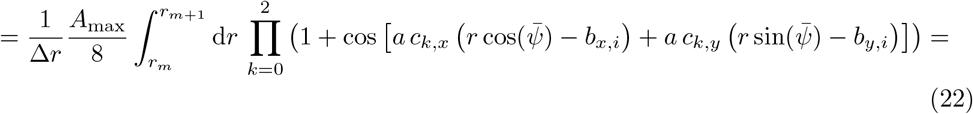

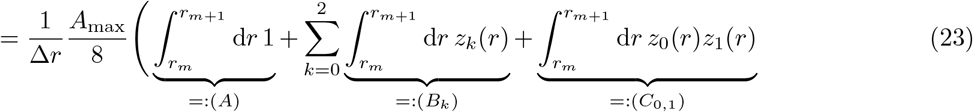

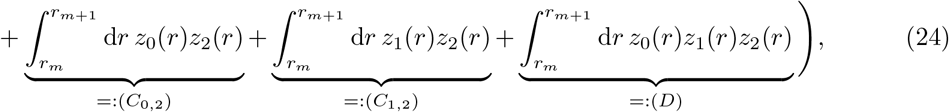

where 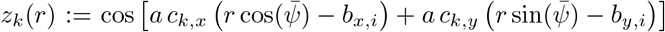 The four parts (A) – (D) are integrated separately. We obtain

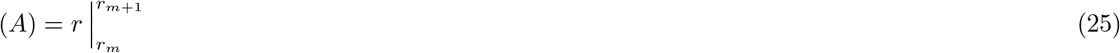

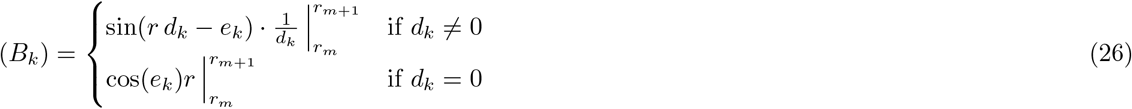

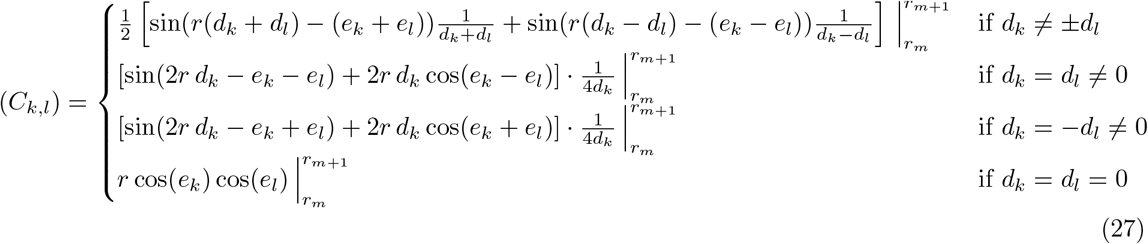

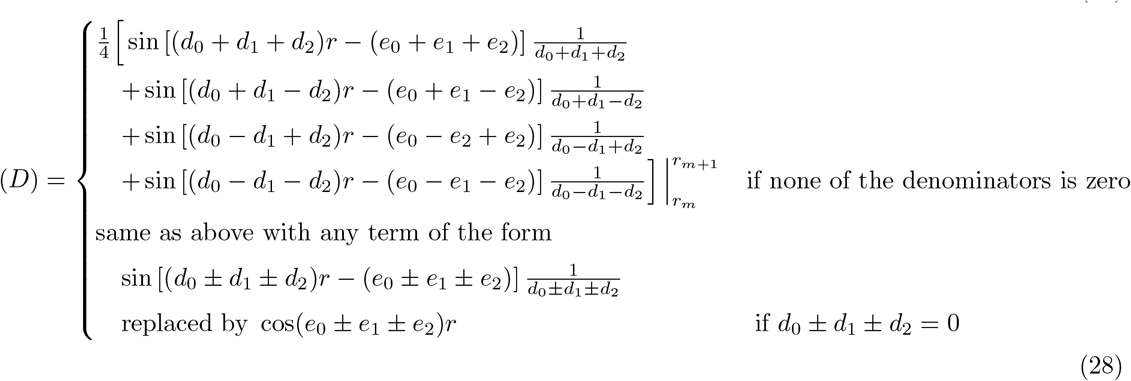

with the abbreviations

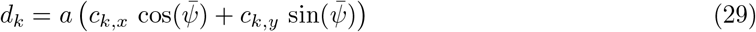

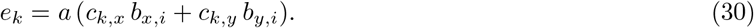

Note that *e*_*k*_ actually depends on the cell index *i* which is omitted in Eqs. (25) - (28) in order to keep the notation simpler.

##### Head-direction tuning

For a conjunctive grid by head-direction cell, the head-direction tuning depends only on the angle (which is fixed when integrating along a straight line through zero) and not at all on the distance from zero. The mean activity is thus obtained from the mean activity of a grid cell without head-direction tuning by multiplying it with *h*(*θ*) defined in Eq. (4):

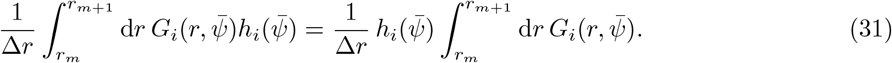

##### Many cells

For more than one cell, the total mean activity (population rate) can simply be calculated as the sum of the mean activities of the single cells

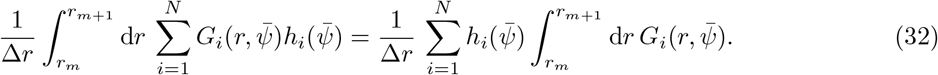

##### Trajectories

The derived analytical description of the mean activity can be applied only to pieces of linear trajectories through zero. For the star-like walk, we can simply integrate

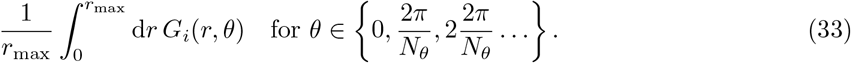

Piecewise linear trajectories and random walk trajectories consist of segments of straight lines that do not necessarily pass through zero. In order to integrate along the *m*-th segment of a trajectory from 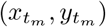 to 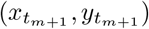 with movement direction *θ*_*m*_, we shift this path segment to the origin by subtracting 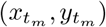 from the grid offset (*x*_off_, *y*_off_) and then integrating

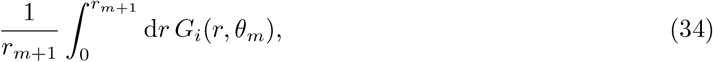

where 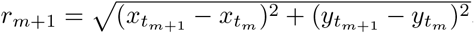.

#### 5.4.2 Upper bound for hexasymmetry of path trajectories

In the following we derive approximations for the expected value of the hexasymmetry 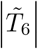 of a path trajectory described by the number of time steps *M*, the movement tortuosity *σ*_*θ*_, and the time step size Δ*t*. These approximations can be used to assess the contribution of the underlying trajectory to the overall hexasymmetry of neural activity.

From the definition of the Fourier coefficients in Eq. (11) and the movement direction distribution in Eq. (**??**), we get the sixth Fourier coefficient of a trajectory:

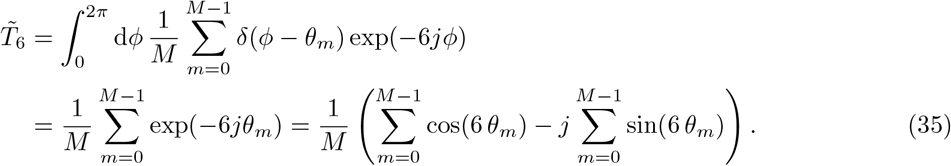

The hexasymmetry of the trajectory can thus be expressed as

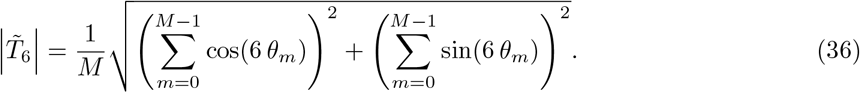

We simplify the sum of squares in Eq. (36)

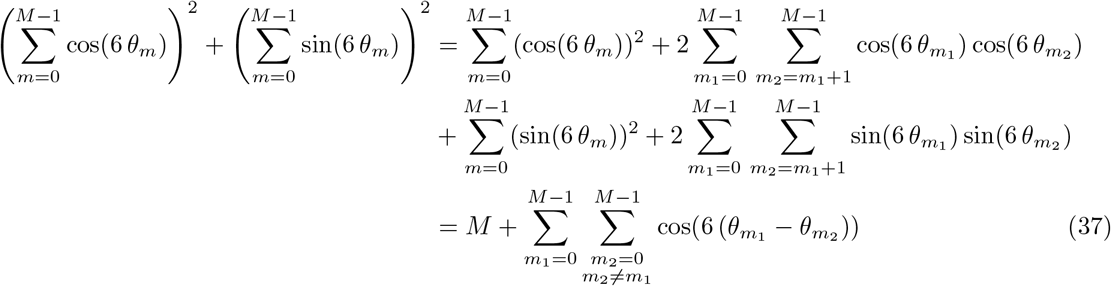

with the help of the multinomial theorem and the trigonometric identity cos(*α − β*) = cos(*α*) cos(*β*) + sin(*α*) sin(*β*). If 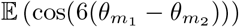 is known, we can compute the expected value of the square of the hexasymmetry of the path trajectory as

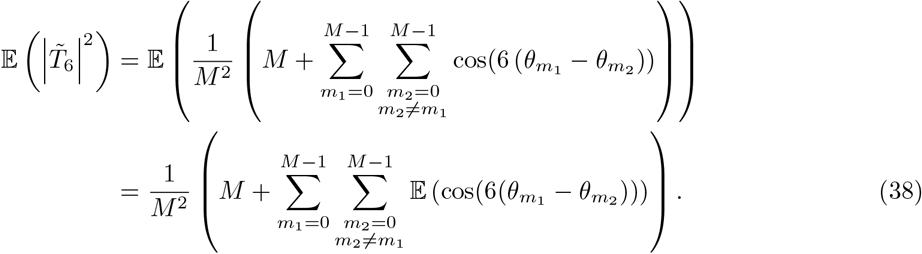

As 𝔼(*X*^2^) = (𝔼(*X*))^2^ + Var(*X*) for any random variable *X*, we can use the result in (38) to obtain the upper bound

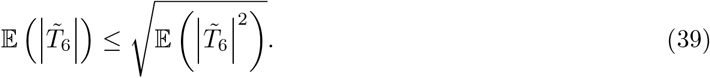

In the following, we focus on the derivation of 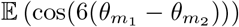. Using the movement statistics

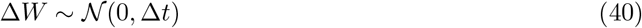

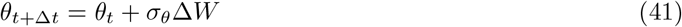

that were introduced below Eq. (1), the distribution of *θ*_*m*_ (after *m* time steps) can be derived when we start at some angle *θ*_0_:

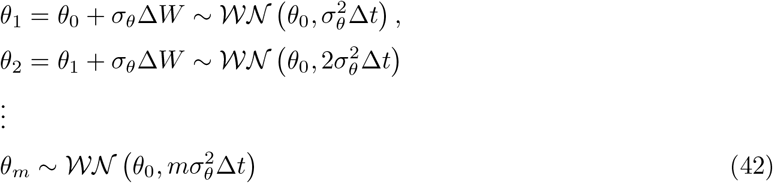

where *𝒲𝒩* (*µ, σ*^2^) denotes the wrapped normal distribution with parameters *µ* and *σ*^2^, which correspond to the mean and variance of the corresponding unwrapped distribution (Jammalamadaka and Sengupta, 2001).

In the following, we will derive the probability distribution of 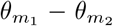 We first define 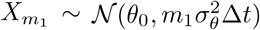 and 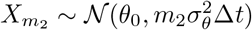 as the unwrapped versions of 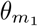 and 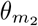, respectively, and 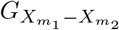 as the distribution function of their difference 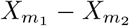. Then, the distribution function (Fisher, 1995) of 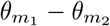 reads as

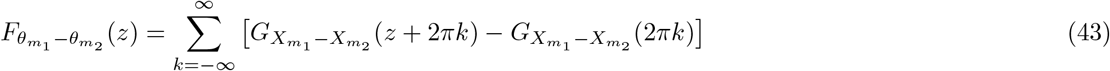

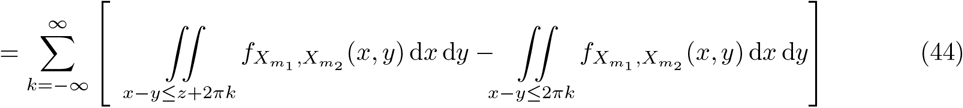

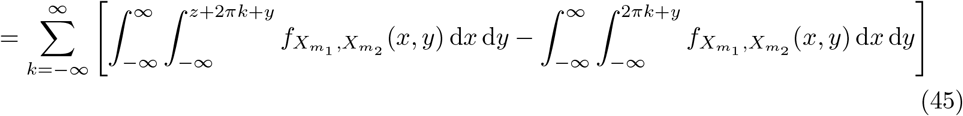

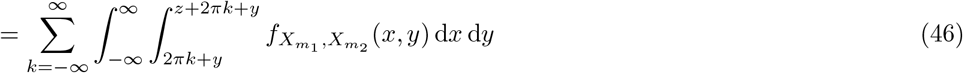

where 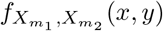 is the joint probability distribution.

In order to calculate the distribution of 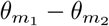, we have to take the dependence between the two angles 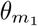 and 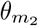 into account. We will first consider the case *m*_1_ *> m*_2_. In this case, the conditional distribution of 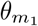 given 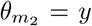 is wrapped normal with conditional mean 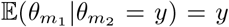 and conditional variance 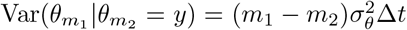 The same can be said about the unwrapped versions of 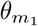 and 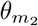. We will use this result to calculate the joint probability distribution

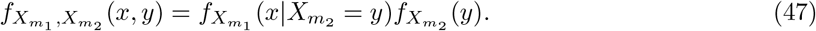

By applying the Leibniz integral rule in (*∗*) we obtain the probability density of 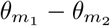

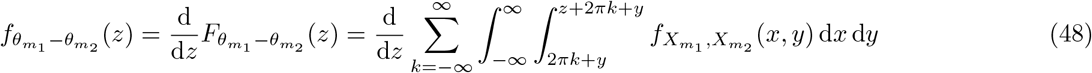

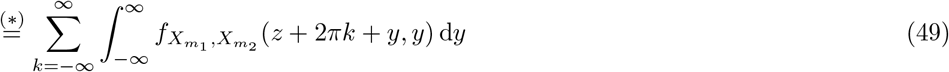

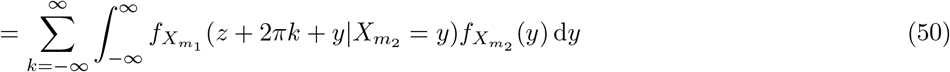

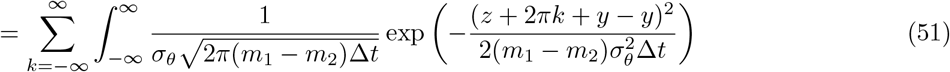

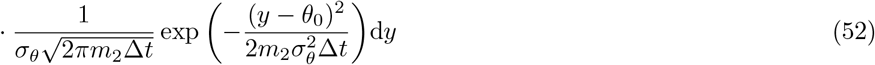

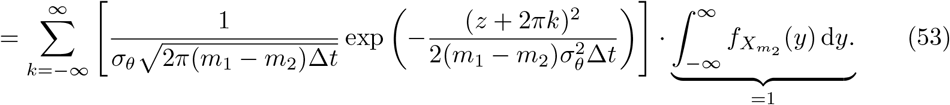

For *m*_2_ *> m*_1_, we derive in the same way as above

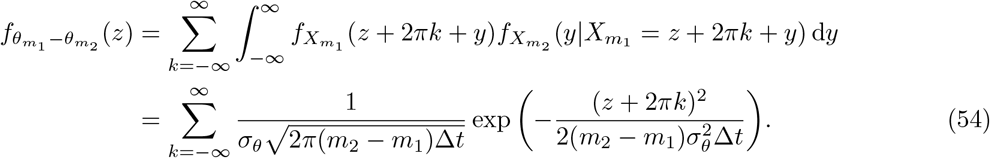

Finally, for *m*_2_ = *m*_1_, we have 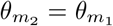. Altogether, we thus get

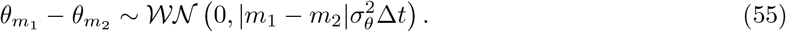

In order to calculate an upper bound for the average path hexasymmetry, we will now use Eq. (55) in Eq. (38). Since 𝔼 (ℛ (*Z*)) = ℛ (𝔼 (*Z*)) for a complex random variable *Z* and the *n*-th moment of a wrapped normal distribution with parameters *µ* and *σ*^2^ is 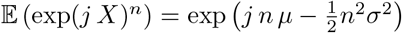, we can derive

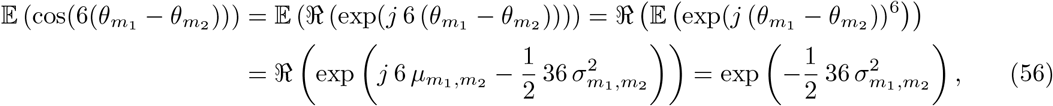

where 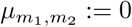 and 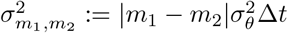 We thus obtain

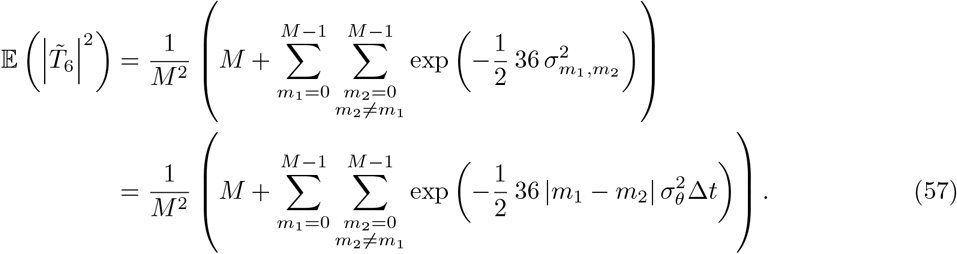

The solid lines in Figure S1 show the square root of Eq. (57) (cf. Eq. (39)).

From Eq. (57) we can derive simplified approximations for two limiting cases. For convenience, we use 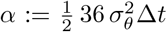 First, if *α »* 1, i.e. if the new direction after one step is almost uniformly distributed or independent of the previous direction, we can neglect the double sum and we have

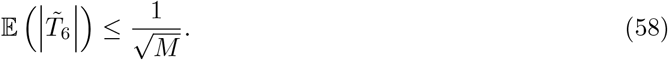

The corresponding line is shown in red in Figure S1. Note that in Figure S1 (with simulation step size Δ*t* = 0.01 s), *α »* 1 corresponds to *σ*_*θ*_ *»* 2.36. A comparison with results from numerical simulations shows that for any *σ*_*θ*_ *>* 3.5 the Eq. (58) constitutes a viable upper bound of the mean path hexasymmetry.

Second, we assume *Mα »* 1, i.e. the direction after *M* steps is almost independent from the original direction. The double sum in Eq. (57) can then be approximated:

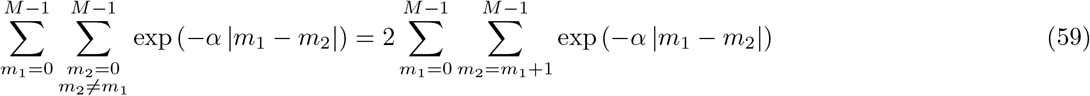

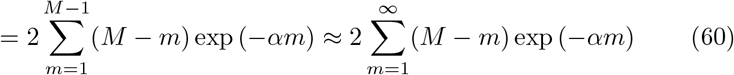

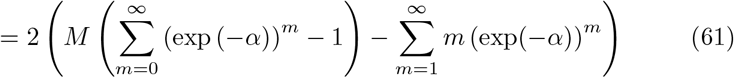

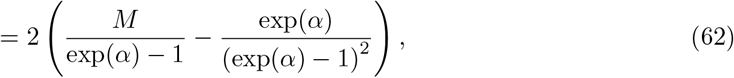

where the first series in (61) is a geometric series and the second series is the polylogarithm function of order -1. We further approximate

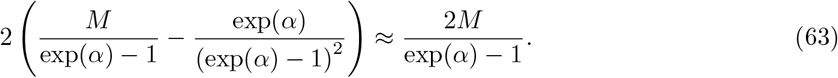

For *α «* 1, the error in the last approximation is 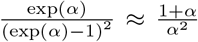, which, if *Mα »* 1, is negligible compared to the remaining term 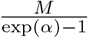 Since 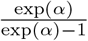 is a strictly monotonically decreasing function of *α*, this approximation does not only hold for *α «* 1 but is good in general. Inserting Eq. (63) into Eq. (57) gives

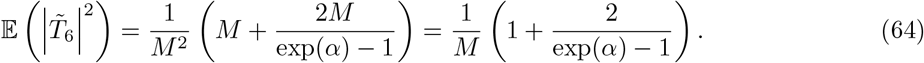

Hence, we get the approximation (for *Mα »* 1)

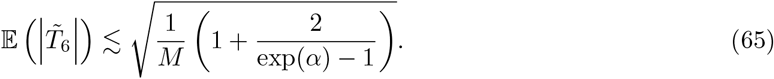

This expression is used to compute the dashed lines in Figure S1, which all have slope 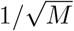 but a prefactor that depends on 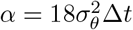.

For *α «* 1 (but still *Mα »* 1), we can use in Eq. (64) the first-order Taylor expansion of the exponential function at 0 to obtain

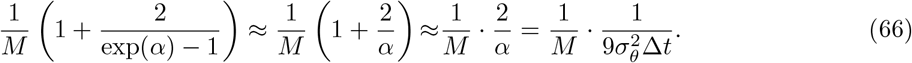

Hence the path hexasymmetry for *α «* 1 and *Mα »* 1 can be approximated by

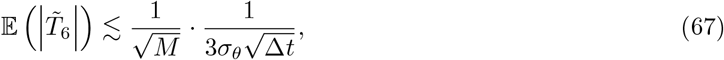

which allows us to see how the key variables *M, σ*_*θ*_, and Δ*t* interact in this limiting case. For instance, given a certain trajectory *A* with *M*_*A*_ steps and random walk parameters *σ*_*θA*_ and Δ*t*_*A*_, we can use Eq. (67) to derive how many steps *M*_*B*_ are necessary in a second trajectory *B* with parameters *σ*_*θB*_ and Δ*t*_*B*_ to achieve the same mean path hexasymmetry. From Eq. (67), we know that the two path hexasymmetries will have the same upper bound if

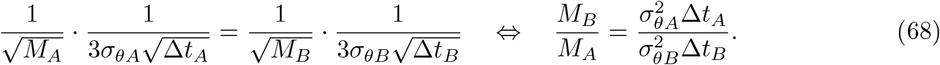

Hence, the number of time steps *M*_*A*_ has to be multiplied by a factor 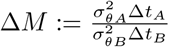

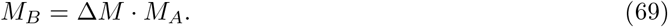

We illustrate the above considerations with an example: Let *σ*_*θA*_ = 1 rad/s^1/2^, *σ*_*θB*_ = 0.5 rad/s^1/2^ and Δ*t*_*A*_ = Δ*t*_*B*_ = 0.01 s. From the given values, we obtain

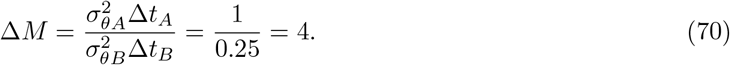

Thus, the mean hexasymmetry value of a trajectory with *σ*_*θA*_ = 1 after *M*_*A*_ time steps is the same as the mean hexasymmetry value of a trajectory with *σ*_*θB*_ = 0.5 after *M*_*B*_ = 4 *· M*_*A*_ time steps. Results from numerical simulations of path hexasymmetries, shown in Figure S1, support the derived theoretical approximations.

### 5.5 Random-field simulations

To quantitatively evaluate the structure-function mapping hypothesis, we set out to simulate a set of grid cells in three-dimensional anatomical space. The grid phases associated with these grid cells follow the correlation structure suggested by (Gu et al., 2018) and (Heys et al., 2014). Our aim is to quantify the clustering of grid phases for a realistically-sized fMRI voxel given this correlation structure.

We use a three-dimensional representation of a voxel with a volume of (3 mm)^3^. Within this voxel, we define a grating of 200^3^ potential grid cells that are equally spaced in the voxel, with a distance between neighbouring cells of 15 *µ*m along the axes of the grating. To generate a set of random but spatially correlated grid phases on this area, we first define two random unit vectors in the complex plane, ***Z***_1_ and ***Z***_2_, for each of the 200^3^ potential grid cells in the voxel; angles of the unit vectors are thus drawn from a uniform distribution on the interval [0, 2*π*). ***Z***_1_ and ***Z***_2_ are further resolved into their real and imaginary components Re(***Z***_*i*_) and Im(***Z***_*i*_), respectively, where *i ∈ {*1, 2*}*. To generate correlations between grid phases, we then convolve the two resulting gratings of 200^3^ components separately with either a Gaussian kernel (Fig. 4K) or a grid kernel (Fig. 4O) to yield the convolved components Re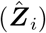 and Im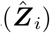. The grid phases can be obtained by first calculating the angles of the new set of complex numbers and normalizing the result by 2*π*:

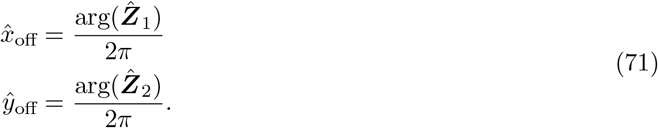

We note that 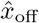 and *ŷ*_off_ are defined on the interval [0, 1) and correspond to the grid phases of a single grid cell mapped to the unit square. Transforming the result to the unit rhombus yields the grid phases *x*_off_ and *y*_off_ in the *x* and *y* direction respectively:

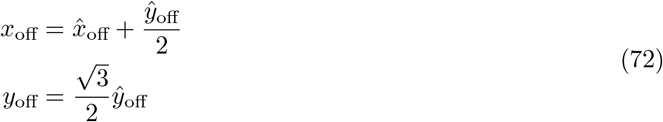

To find the average pairwise grid phase distances as a function of the pairwise anatomical distances, 10^8^ pairs of grid cells are sampled randomly from the uniform distribution defined on the discrete space of grating cell positions. The Euclidean distance in anatomical space between the two grid cells in each pair is calculated and sorted into 50 bins of equal width on the interval [10, 500] *µ*m. Then, for each pair of grid cells, *n*_1_ and *n*_2_, 8 copies of the grid phase (*x*_off,2_, *y*_off,2_) of the second cell *n*_2_ are made, which are offset from the initial position of the grid phase such that they are positioned at the same phase within unit rhombi laid end-to-end on a 3 × 3 grid. The minimum distance between the grid phase of the cell *n*_1_ and the grid phase of each of the copies of the cell *n*_2_ is taken as the pairwise phase distance. Finally, the pairwise distance between grid phase offsets per distance bin is obtained by taking the mean over all grid cell pairs whose Euclidean distance in anatomical space falls into the corresponding bin (Fig. 4H, L, P).

To estimate the clustering concentration parameter *κ*_*s*_ in Fig. 4, the phases 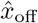 and *ŷ*_off_ are mapped to a circular distribution by multiplying them with 2*π*. The sets of grid phases 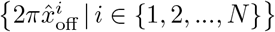 and 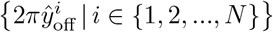 are then each separately fit to a one-dimensional von Mises distribution to obtain a clustering concentration parameter for each axis. The final value of *κ*_*s*_ is taken as the average of these two values.

### 5.6 Parameter estimation

The hexasymmetry values of the tested hypotheses depend on the respective parameters. We investigate two scenarios for each hypothesis: an “ideal” set of parameters that results in a large (basically maximal) hexasymmetry value and a “realistic” set of parameters that we try to derive from experimental data. All sets of parameters are summarized in Table 1. We comment here on the “realistic” sets of parameters and how we derived them:

#### Conjunctive grid by head-direction hypothesis

For the “realistic” value of *σ*_*c*_ = 3°, we use the 95% confidence interval of 12.4° from the third row of the table in the Supplementary Figure 5b of Doeller et al. (2010) and translate this value to a Gaussian standard deviation, assuming a 95% confidence interval of 4*σ*_*c*_.

For the “realistic” value of *κ*_*c*_ = 4, we refer to the Supplementary Figure 3 of Doeller et al. (2010). Visually, the tuning curves usually cover between one sixth and one third of the circle. Stronger head-direction tuning contributes the most to the resulting hexasymmetry, so we choose a value of *κ*_*c*_ = 4, which corresponds to a tuning width of roughly one sixth of the circle. For further discussion on *κ*_*c*_, see also Supplementary Figure Fig. S9.

The “realistic” value of *p*_*c*_ = 1/3 was chosen based on the values reported by Sargolini et al. (2006) and Boccara et al. (2010). In our simulations, we observed that the hexasymmetry shows a linear dependence on the fraction of conjunctive cells up to a noise floor (Fig. S10).

#### Repetition suppression hypothesis

To obtain “realistic” parameter values, we simply divide the “ideal” ones by two, which results in a “realistic” adaptation time constant *τ*_*r*_ = 1.5 s and a “realistic” adaptation weight *w*_*r*_ = 0.5. “Ideal” parameter values attenuate the firing rate by *≈* 20% (Fig. 3A, with 24% when running aligned with the grid axes and 18% when running misaligned to the grid axes). “Realistic” parameter values attenuate firing rates to *≈* 16% (for a dependence of hexasymmetry on the two parameters, see Fig. 3B). Such a decrease is similar in magnitude to the attenuation of 5% of firing rates in Reber et al. (2023) (their Figure 2B, right). The value of the “realistic” time constant is also comparable to the time scales found in two electrophysiological studies: firstly, Giocomo and Hasselmo (2008) showed that the time constant in the slow component of the hyperpolarization-activated current (*I*_*h*_) in the ventral mEC is 2–3 s (their Figure 2F). Secondly, Magistretti and Alonso (1999) found a second-long inactivation of a persistent sodium current in EC layer II cells.

#### Structure-function mapping hypothesis

The “realistic” value of the concentration parameter for clustering, *κ*_*s*_ = 0.1, is derived from the random-field simulations in Fig. 4G–Q and a comparison to results in Gu et al. (2018) and Heys et al. (2014). Figure 4G in Gu et al. (2018) shows for grid cells in the rat mEC the relationship between their pairwise physical distance and their pairwise phase distance. Similarly, for grid cells in the mouse mEC, Heys et al. (2014) in their Figure 4B show the correlation of (one-dimensional) grid firing maps as a function of anatomical pairwise distance between grid cells. Together, these data serve as evidence for more similar grid cell firing for anatomically close grid cells (distance *<* 100 *µ*m) — with decreasing similarity for increasing anatomical distance. In our random-field simulations (Fig. 4), we mimic these experimental results by choosing the width of a spatial correlation kernel accordingly (Fig. 4K, Gaussian kernel with standard deviation: 30 *µ*m). Consequently, in the simulations, the initial rise of phase distance as a function of anatomical distance (Fig. 4L) reflects the experimental curves. When grid cells in anatomical space are further apart than roughly 100 *µ*m, the behavior of the experimental curves by Gu et al. and Heys et al. is less consistent. In Heys et al.’s Figure 4B there is a slight trough in grid-firing correlation at about 100—150 *µ*m (corresponding to a slight peak in grid-phase distance at physical distance 200 *µ*m in Gu et al.’s Figure 4G), followed by an increase in grid-firing similarity (decrease of phase distance) for anatomical distances of about *>* 150 *µ*m (*≈* 300 *µ*m). In our simulations, we reflect such a non-monotonous behavior by introducing a grid-like correlation kernel (Fig. 4O) with a grid spacing of 300 *µ*m. Applying this grid-like correlation kernel to the random field in Fig. 4G results in a phase-distance curve with a slight dip around 300 *µ*m (Fig. 4P), just like in the experimental curves in Gu et al. (2018). For both cases, the Gaussian kernel and the grid-like kernel, we find that the resulting phase clustering for an anatomical voxel of realistic size is quite low, i.e. *κ*_*s*_ is well below 0.1 (Fig. 4M,Q). We use 0.1 as a conservative estimate of the “realistic” value for *κ*_*s*_ in our further simulations. For the central phase of the cluster, we always use (without loss of generality) the origin: *µ*_*s*_ = (0, 0).

### 5.7 Implementation of previously used metrics

We applied three previously used metrics to our framework: the GLM method by Doeller et al., 2010; the GLM method with binning by Kunz et al., 2015; and the circular-linear correlation method by Maidenbaum et al., 2018 (Fig. S8).

In brief, in the GLM method (e.g. used in Doeller et al., 2010), the hexasymmetry is found in two steps: the orientation of the hexadirectional modulation is first estimated on the first half of the data by using the regressors *β*_1_ cos(6*θ*_*t*_) and *β*_2_ sin(6*θ*_*t*_) on the time-discrete fMRI activity (Eq. 9), with *θ*_*t*_ being the movement direction of the subject in time step *t*. The amplitude of the signal is then estimated on the second half of the data using the single regressor *β* cos[6(*θ*_*t*_ *− ϕ*)], where *ϕ* = arctan(*β*_2_*/β*_1_)/6. The hexasymmetry is then evaluated as *H* = *β/*2.

The GLM method with binning (e.g. used in Kunz et al., 2015) uses the same procedure as the GLM method for estimating the grid orientation in the first half of the data, but the amplitude is estimated differently on the second half by a regressor that has a value 1 if *θ*_*t*_ is aligned with a peak of the hexadirectional modulation (aligned if |*θ*_*t*_ *− ϕ*| % 60° *<* 15°; %, modulo operator) and a value of *−*1 if *θ*_*t*_ is misaligned. The hexasymmetry is then calculated from the amplitude in the same way as the GLM method.

The circular-linear correlation method (e.g. used in Maidenbaum et al., 2018) is similar to the GLM method in that it uses the regressors *β*_1_ cos(6*θ*_*t*_) and *β*_2_ sin(6*θ*_*t*_) on the time-discrete mean activity, but instead of using *β*_1_ and *β*_2_ to estimate the orientation of the hexadirectional modulation, the beta values are directly used to estimate the hexasymmetry using the relation 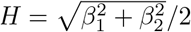.

### 5.8 Tuning widths and the large *κ* approximation

To provide a comparison between our tuning widths (using the small angle approximation 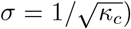 with the breadth of directional tuning used in Sargolini et al. (2006), we simulated random walk trajectories of length *T* = 9000 s, time step *dt* = 0.01 s, tortuosity parameter *σ*_*θ*_ = 0.5 rad/s^2^ and a speed of 10 cm/s in an infinite environment. We then calculated the mean firing rate in angular bins of size 6° from 0° to 360° using Eq. 3 (the conjunctive grid by head-direction cell hypothesis) for varying values of the tuning width *κ*_*c*_. From the mean firing rates, we calculate the length *ρ* of the resultant vector by vector summation across all 60 bins and then dividing by the sum of individual vector lengths. The breadth of directional tuning is expressed as the angular standard deviation using the equation 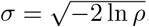 To connect this result to the concentration parameter of a von Mises distribution, we additionally used an approach where we calculate the resultant length using a ratio of modified Bessel functions of the first kind such that 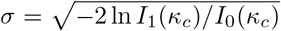 where *I*_*n*_(*κ*_*c*_) is the modified Bessel function of the first kind of order ‘n’. The results for this comparison are summarized in Fig. 9.

To quantify the hexasymmetry when using tuning widths reported in empirical studies (Doeller et al., 2010; Sargolini et al., 2006), we simulated 50 instances of the conjunctive grid by head-direction cell hypothesis for the tuning widths 8° (corresponding to our “ideal” parameters), 29° (corresponding to Doeller et al.), and 55° (corresponding to Sargolini et al.). For a jitter of *σ*_*c*_ *<* 10°, we found significant hexadirectional modulation for a tuning width of 29°, while the hexasymmetry appears to be at noise levels for tuning widths larger than about 35° such as those reported by Sargolini et al., regardless of the value of the jitter. (Fig. S9C). However, it should be noted that unlike in our simulations, the tuning widths reported by these studies are a population average that might consist of a large number of widely tuned cells and a small subpopulation of narrowly tuned cells. In this case, the finely tuned subpopulation would still give rise to a hexadirectionally modulated signal under the framework of the conjunctive grid by head-direction cell hypothesis. Furthermore, Doeller et al. and Sargolini et al. had different criteria for identifying HD cells, which could account for the significantly different tuning widths in these studies.

### 5.9 Numerical simulations

All simulations were implemented in Python 3.10 using the packages NumPy, SciPy, Numba, and JobLib. Matplotlib was used for plotting. Inkscape was used for the final adjustment to the figures.

## Acknowledgements

We would like to thank Tiziano D’Albis for discussions.

This work was funded by the German Research Foundation (DFG, Project number 327654276 – SFB 1315 to RK), the German Federal Ministry of Education and Research (01GQ1705 to RK), and the Einstein Foundation Berlin (to IK). LK received funding from the German Research Foundation (DFG; project nos. 447634521 and 527084865), the return program of the Ministry of Culture and Science of the State of North Rhine-Westphalia, the Federal Ministry of Education and Research (BMBF; 01GQ1705A), and by NIH/NINDS grant U01 NS113198-01.

## Author contributions

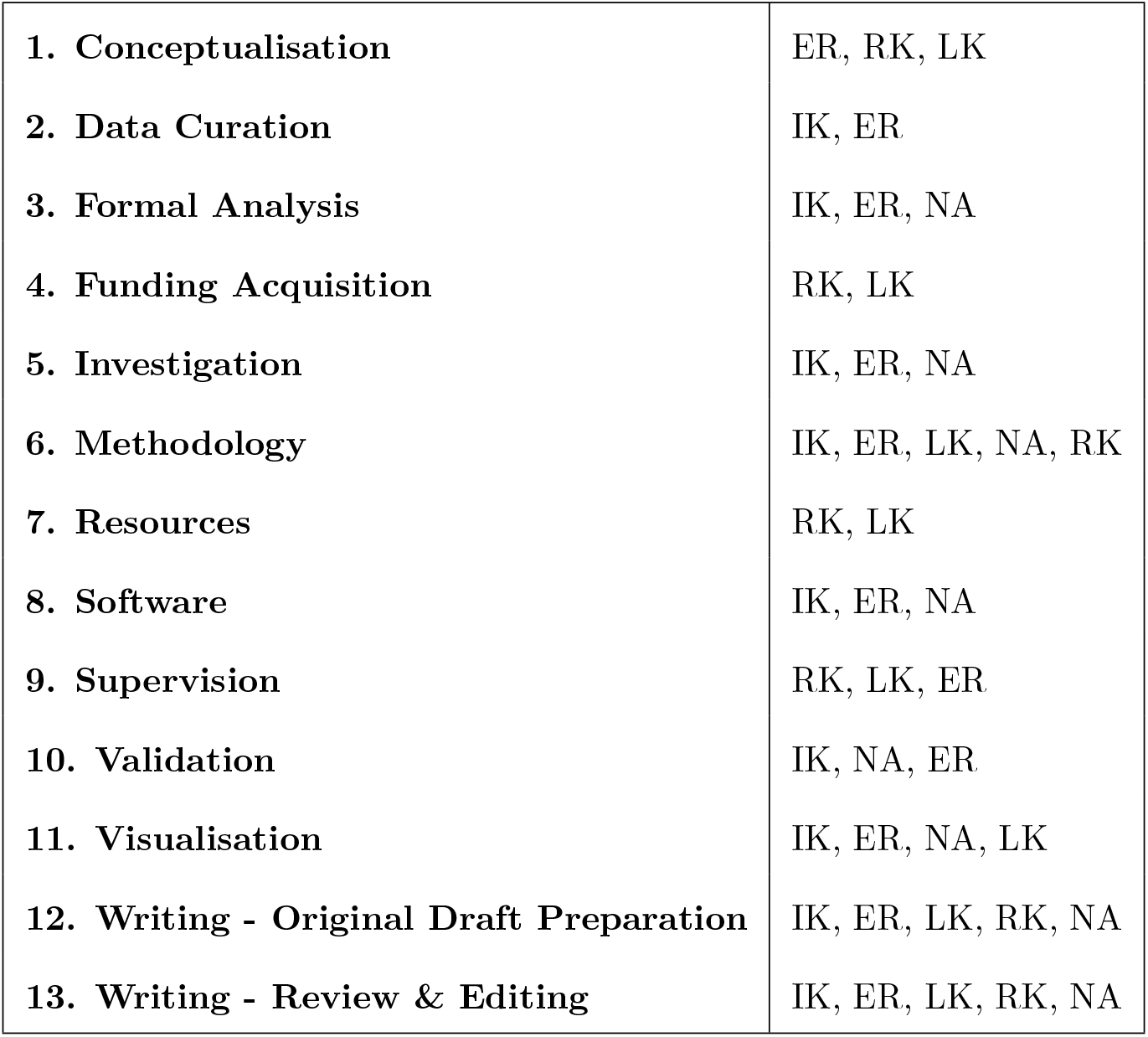

## Data and code availability

The code used to generate the data is available at https://github.com/ikhwankhalid/grid_bold.

---

**Figure S1:**
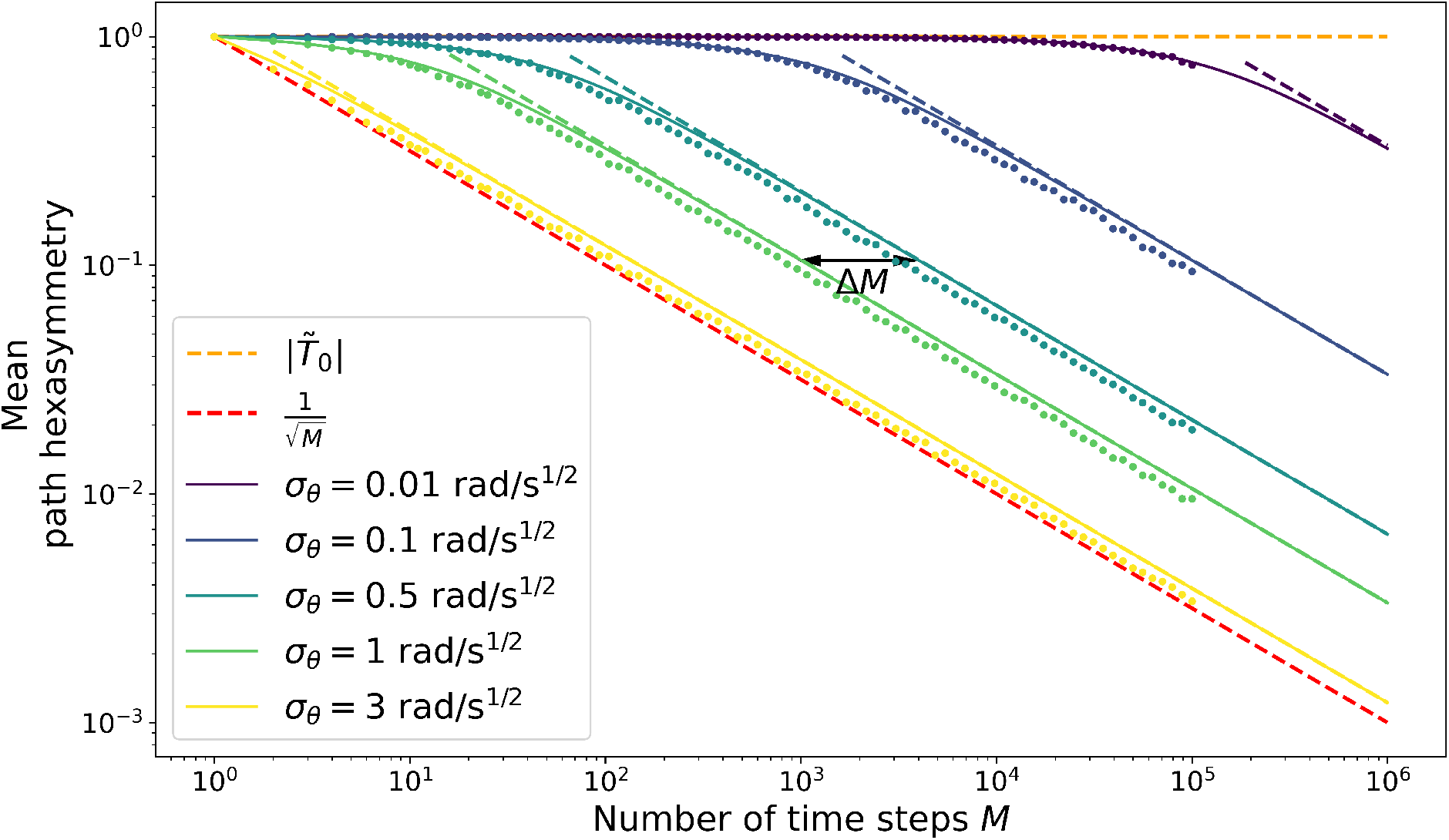
Hexasymmetry of random walk trajectories. The (horizontal) orange dashed line shows the offset (or maximum) of the path hexa symmetry, 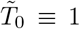. The (diagonal) red dashed line shows the average path hexasymmetry 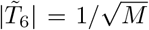 associated with randomly sampling a movement direction at each time step from a uniform distribution on the interval [0, 2*π*). Solid colored curves show an upper bound for the mean path hexasymmetry of a random walk as a function of the number of time steps (square root of Eq. (57)) for five different movement tortuosities *σ*_*θ*_ and a simulation time step size Δ*t* = 10 ms. The corresponding five colored dashed lines show an approximation 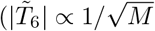 Eq. (65)) to the solid curves; the approximation is excellent if the number *M* of time steps is large enough. Colored dots show the respective mean path hexasymmetries obtained from numerical simulations (Eq. (1)). The black arrow shows the multiplicatory shift in the number of time steps that is necessary to obtain the same hexasymmetry for a trajectory with *σ*_*θ*_ = 0.5 rad/s^1/2^ as for a trajectory with *σ*_*θ*_ = 1 rad/s^1/2^, as derived in Eq. (70).

**Figure S2:**
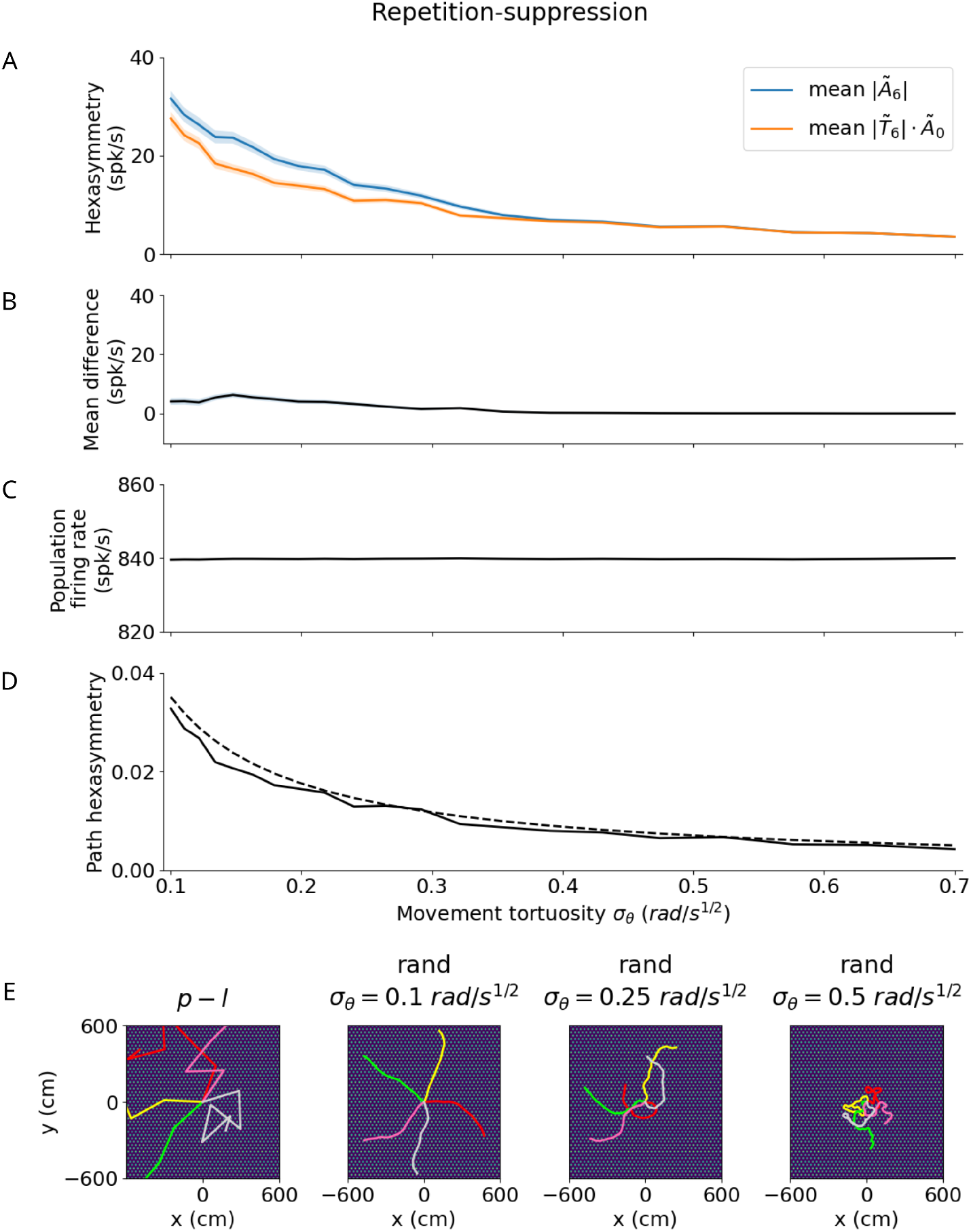
Effect of tortuosity on the hexasymmetry for the repetition-suppression hypothesis with random walks (Δ*t* = 0.01 s, *T* = 9000 s, *τ*_*r*_ = 3 s, *w*_*r*_ = 1). (**A**) The hexasymmetry *Ã*_6_ (blue) and the scaled path hexasymmetry 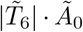 (orange) as a function of the movement tortuosity *σ*_*θ*_ of a random walk. The shaded areas represent the standard error when averaging over 100 realizations of the trajectory, each with an initial direction sampled from a random uniform distribution on the interval [0, 2*π*). (**B**) The mean difference between |*Ã*_6_| and 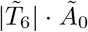 as a function of the movement tortuosity *σ*_*θ*_. Note that 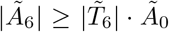 for *σ*_*θ*_ ≤ 0.4. For *σ*_*θ*_ *>* 0.4 the two curves begin to overlap: repetition-suppression ceases to have a significant effect and the hexasymmetry |*Ã*_6_| is primarily dictated by the *∼* path hexasymmetry. The dip in the mean difference near *σ*_*θ*_ ∼ 0.1 is thought to be due to numerical noise. (**C**) The mean firing rate *Ã*_0_ does not depend on movement tortuosity *σ*_*θ*_, indicating that any effect of repetition-suppression on the population firing rate is small. (**D**) Path hexasymmetry 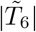 as a function of the movement tortuosity *σ*_*θ*_. The solid line depicts the mean path hexasymmetry averaged over 100 realizations of a trajectory, while the dashed line plots the analytical result in Eq. (65). (**E**) Leftmost panel: Five examples (colored line segments) of a piecewise linear (“p-l”) trajectory; each linear path segment has a length of 300 cm. Three rightmost panels: Five example random walk (“rand”) trajectories (colored curves) for three different values of the movement tortuosity *σ*_*θ*_ with total simulation time *T* = 60 s and a total path length of 600 cm. For illustration purposes, the initial directions of example random walk trajectories were chosen such that they are regularly distributed on the interval [0, 2*π*). When viewed on a scale within the range of *±* 600 cm, increasing the tortuosity from 0.1 to 0.5 results in increasingly more curved trajectories. The bright dots in all panels of (E) show the grid fields with grid spacing 30 cm, which is small when viewed on this scale. In the simulations for Figs. 2–5, the random walk trajectories have total lengths of 90, 000 cm or longer and use a movement tortuosity of 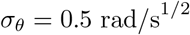.

**Figure S3:**
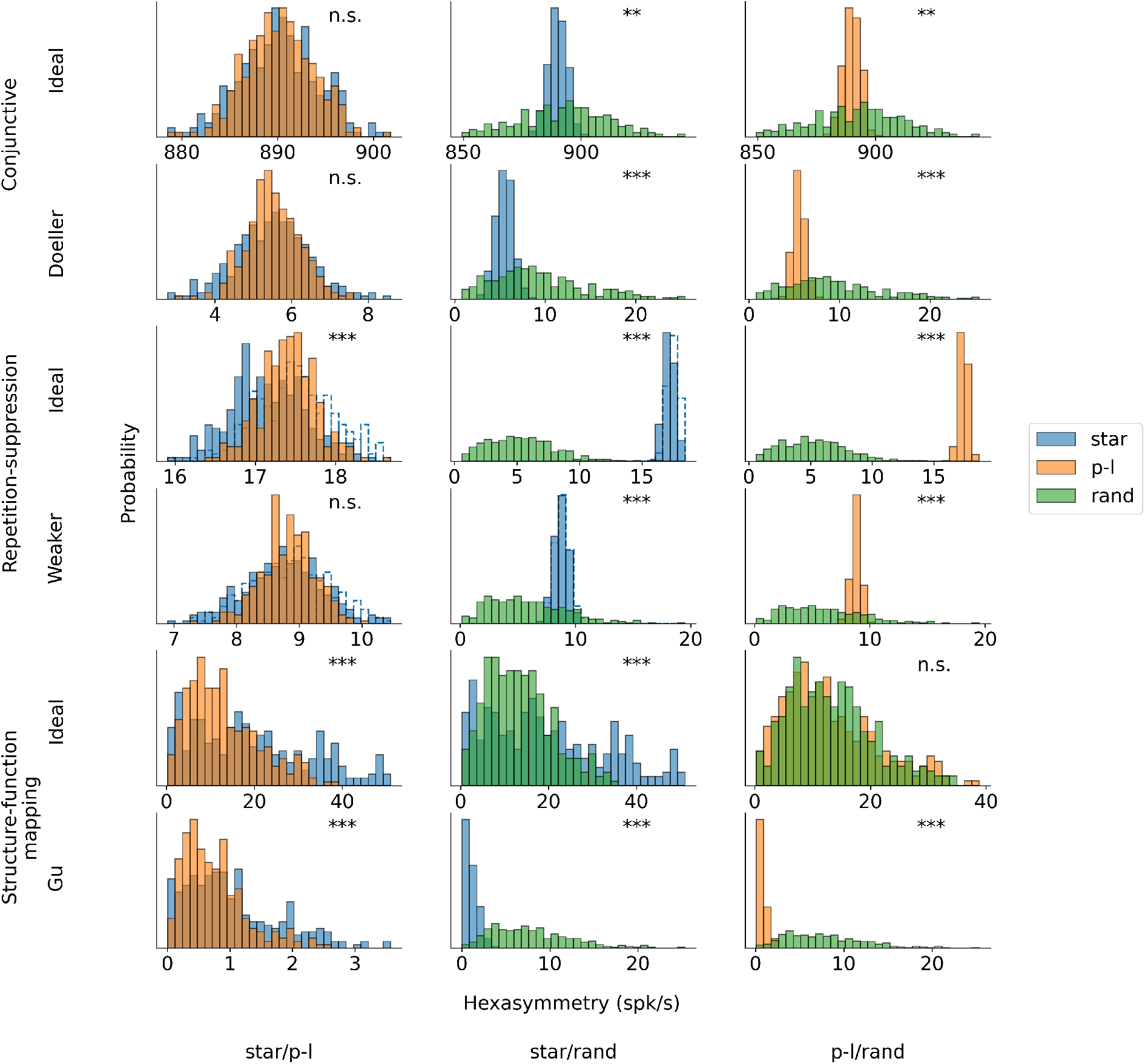
Pair-wise comparisons of the hexasymmetry values from different trajectory types for each set of parameters. The compared trajectory types are star-like walks (“star”), piecewise-linear walks (“p-l”), and random walks (“rand”). For each hypothesis, we calculate the hexasymmetry for ideal parameters (conjunctive: *p*_*c*_ = 1, *κ*_*c*_ = 50 rad^*−*2^, *σ*_*c*_ = 0; repetition suppression: *τ*_*r*_ = 3 s, *w*_*r*_ = 1; clustering: *κ*_*s*_ = 10) as well as more realistic parameters (conjunctive: *p*_*c*_ = 0.33, *κ*_*c*_ = 4 rad^*−*2^, *σ*_*c*_ = 3°; repetition suppression: *τ*_*r*_ = 1.5 s, *w*_*r*_ = 0.5; clustering: *κ*_*s*_ = 0.1). In the case of the repetition-suppression hypothesis, the solid blue bars show the star-like walk with a carry-over of the repetition-suppression mechanism when teleporting between different path segments, while the transparent bars with the dotted blue borders represent the star-like walk with no carry-over of the repetition-suppression effect across different path segments. For the star-like walk, the starting phase of the star is sampled from a uniform distribution across the unit rhombus between realizations, and remains constant within each realization of the star-like walk trajectory. The direction of movement for both the star-like walk and the piecewise linear walk is sampled randomly without replacement from the integer angles 0, 1, 2, …, 359 °. The parameters for the random-walk scenario are *T* = 9000 s and Δ*t* = 0.01 s. Each hypothesis condition was simulated for 300 realizations. ***, *P <* 0.001; **, *P <* 0.01; n.s., not significant. Note that the scales of the horizontal axes are different across subpanels.

**Figure S4:**
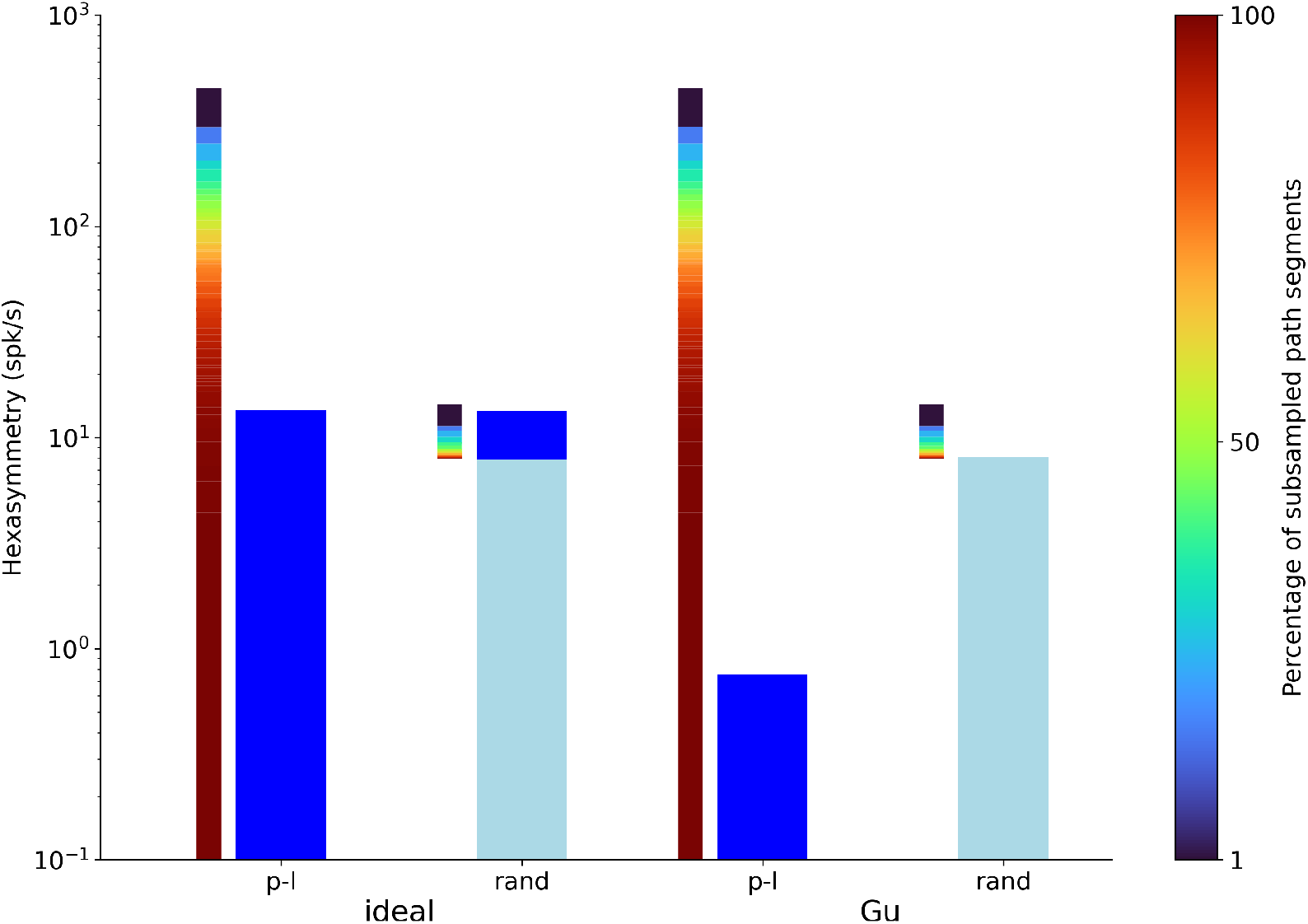
Hexasymmetry resulting from piecewise linear walks (“p-l”) and random walks (“rand”) as a function of the percentage of subsampled path segments for the structure-function mapping hypothesis. Values of “ideal” parameters (left) and more realistic parameter values (“Gu”; right) are identical to the ones used in Fig. 5. The thinner gradient bars show the percentage of subsampled path segments required to produce the corresponding scaled path hexasymmetry 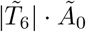 The thicker dark blue and light blue bars represent the hexasymmetry *Ã*_6_ and the path hexasymmetry multiplied by the mean firing rate 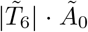, respectively. The percentages of path segments were subsampled from 1% to 100% in steps of 1%. In the case of the random walk trajectories, e.g. subsampling 100% of the path segments yields 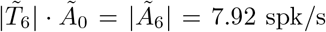 For each percentage value, the path hexasymmetry was averaged over 100 different realizations of the piecewise linear or random walk trajectory.

**Figure S5:**
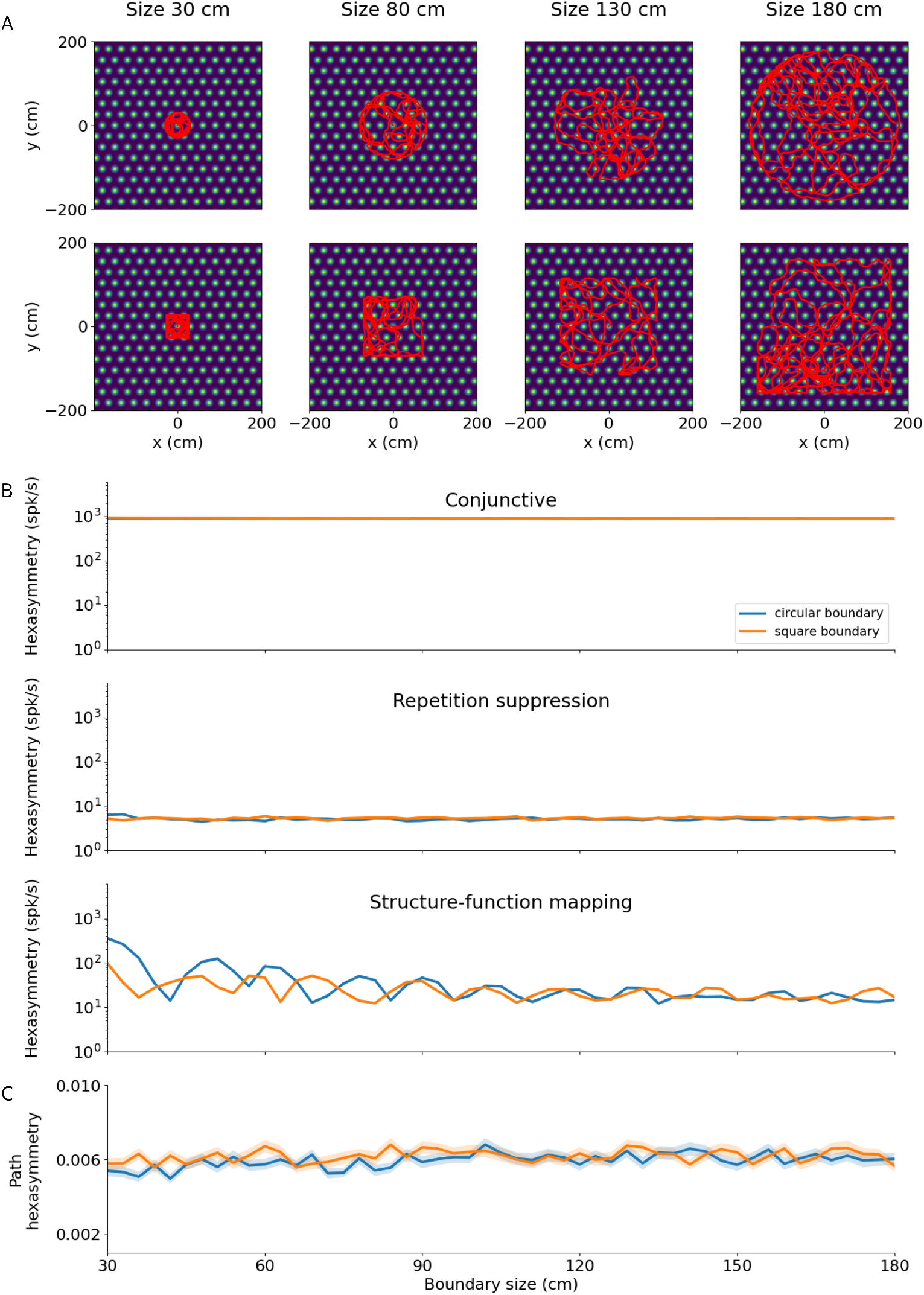
Effect of size and shape of finite environments on hexasymmetry. (**A**) Examples of bounded random walk trajectories (Δ*t* = 0.01 s, *T* = 9000 s, *v* = 10 cm/s, *σ*_*θ*_ = 0.5 rad/s^1/2^; see Section 5.1.2 for details on the generation of bounded trajectories) within square boundaries (bottom; “size” indicates half of side length) or circular boundaries (top; “size” indicates radius) of varying sizes (increasing from left to right). Trajectories are overlaid on the firing field of an example grid cell. The lengths of the depicted trajectories are modified for illustration purposes, and do not reflect the full extent of trajectories used to calculate hexasymmetries. (**B)** Hexasymmetry for the three hypotheses for different sizes of the boundaries; blue, circular boundary; orange, square boundary. For all three hypotheses, the “ideal” parameter sets were used; for values of the parameters, see Table 1. Overall, the obtained values of the hexasymmetry have a weak (if any) dependence on boundary shape and size (apart from fluctuations due to noise in different realizations), and the obtained values are similar in magnitude to those obtained in infinite environments: Fig. 2F for “conjunctive”, Fig. 3E for “repetition suppression”, and Fig. 4C for “structure-function mapping”. **(C)** Path hexasymmetry for different sizes of the boundaries. In (B) and (C), lines represent the mean and shaded areas represent the standard error as obtained from 50 trajectories.

**Figure S6:**
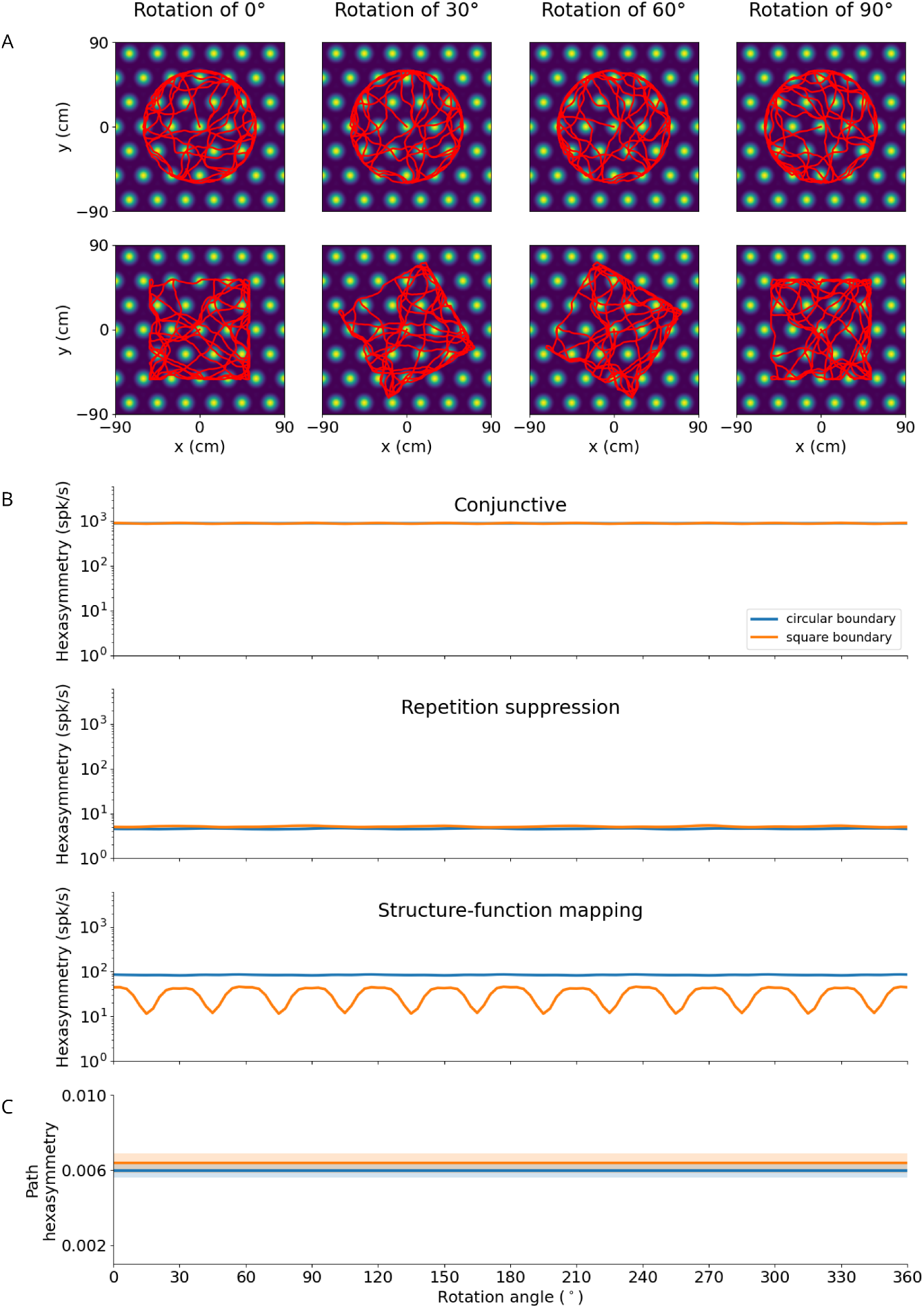
Effect of rotation of finite environments on hexasymmetry. (**A**) Examples of bounded random walk trajectories (Δ*t* = 0.01 s, *T* = 9000 s, *v* = 10 cm/s, *σ*_*θ*_ = 0.5 rad/s^1/2^; see Section 5.1.2 for details on the generation of bounded trajectories) within square boundaries (bottom; “size” indicates half of side length) within circular (top) or square (bottom) boundaries for different rotation angles (numbers at top) relative to the grid orientation. Trajectories are overlaid on the firing field of an example grid cell. The lengths of the depicted trajectories are modified for illustration purposes, and do not reflect the full extent of trajectories used to calculate hexasymmetries. (**B**) Hexasymmetry for the three hypotheses for different rotation angles of the trajectory; blue: circular boundary; orange: square boundary. The periodic fluctuation in neural hexasymmetry for the conjunctive hypothesis with a square environment is due to the alignment of the edges of the square with the grid axes whenever the trajectory is rotated by multiples of 30° and 60° combined with the tendency of the subject to move along the walls of the boundary. With circular boundaries, these fluctuations are closer to the order of the standard error, and are due to a combination of directional bias in the trajectory and noise in the arrangement of grid phase offsets. Otherwise, the obtained values of the hexasymmetry have a weak (if any) dependence on boundary shape and orientation (apart from fluctuations due to noise in different realizations), and the obtained values are similar in magnitude to those obtained in infinite environments: Fig. 2F for “conjunctive”, Fig. 3E for “repetition suppression”, and Fig. 4C for “structure-function mapping”. For all three hypotheses, the “ideal” parameter sets were used; for values of the parameters, see Table 1. (**C**) Path hexasymmetry does not depend on the rotation angle of the trajectory. In (B) and (C), lines represent the mean and shaded areas represent the standard error as obtained from 50 random walk trajectories with the same parameters as in Table 1.

**Figure S7:**
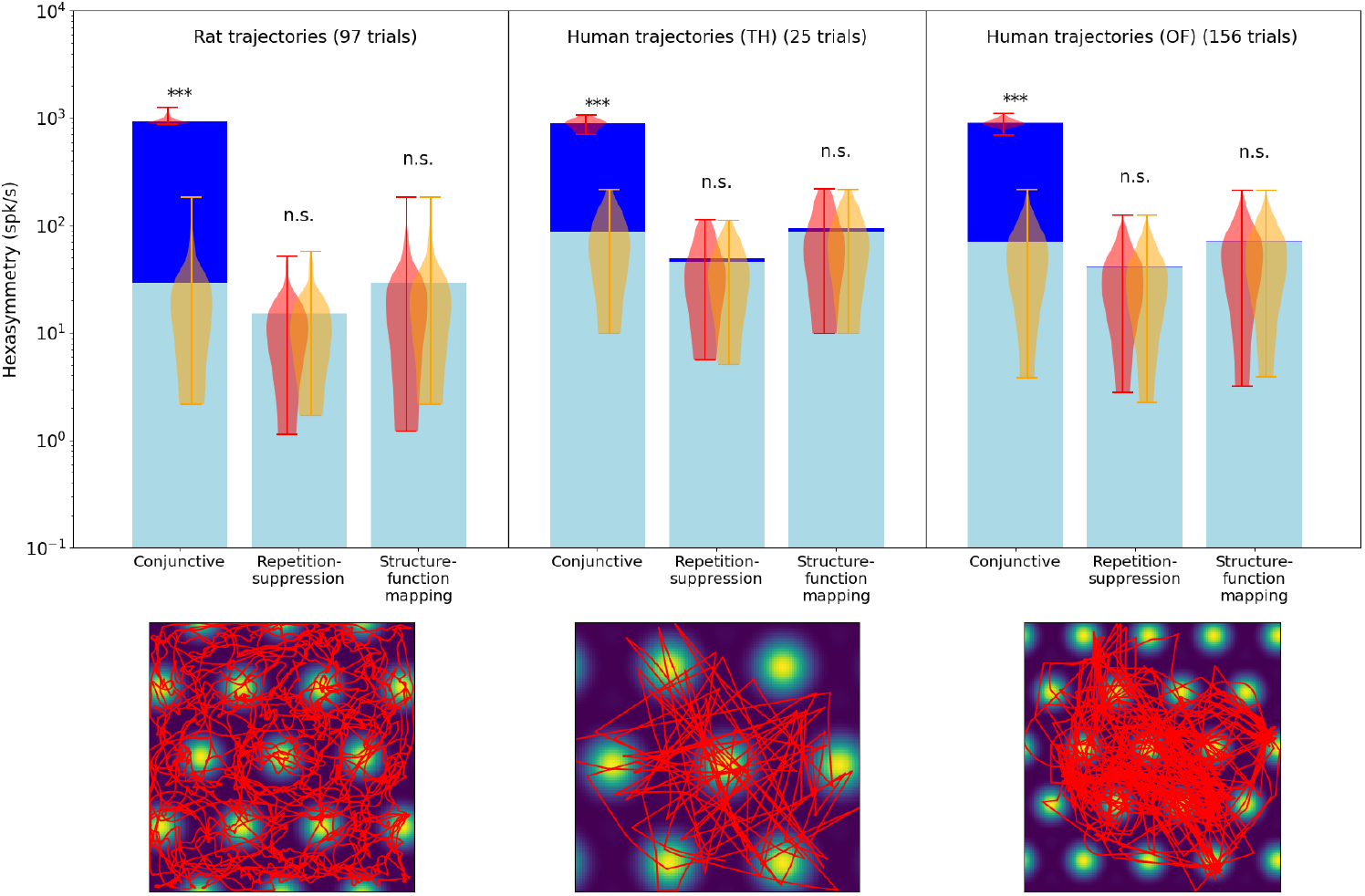
Comparison of the hexasymmetry resulting from the three hypotheses when using trajectories from rats and humans. For each hypothesis condition, we used 97 rat trajectories (Sargolini et al., 2006) and 181 human trajectories across two data sets (Kunz et al., 2015, 2021) to estimate neural hexasymmetry and path hexasymmetry. Example trajectories are overlaid on the firing field of a grid cell (bottom panels). The human trajectories were rescaled such that the subject traversed 3-4 grid firing fields (Hafting et al., 2005) when crossing the environment. For each data set, the corresponding neural hexasymmetry and path hexasymmetry were evaluated. In the top panel we show for each setting the average neural hexasymmetry |*Ã*_6_ |for 1024 cells (dark blue bars) and the average contribution of the trajectory 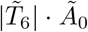 (light blue bars) where *Ã*_0_ is the (variable) average population activity; see Methods for definitions of symbols. The violin plots depict the distributions for firing-rate hexasymmetry (red) and path hexasymmetry (orange). While the human trajectories do not result in significant neural hexasymmetry under the repetition suppression hypothesis in this data, we note that there is a variety of different virtual-reality navigation tasks in humans with varying degrees of straightness. For example, in Horner et al. (2016), the navigation paths very clearly resemble piecewise-linear walks. In such studies, repetition-suppression effects may play a role in the emergence of grid-like representations. For each hypothesis, we calculated the hexasymmetry for “ideal” parameters (conjunctive: *p*_*c*_ = 1, *κ*_*c*_ = 50 rad^*−*2^, *σ*_*c*_ = 0; repetition suppression: *τ*_*r*_ = 3 s, *w*_*r*_ = 1; clustering: *κ*_*s*_ = 10). ***, *P <* 0.001; n.s., not significant.

**Figure S8:**
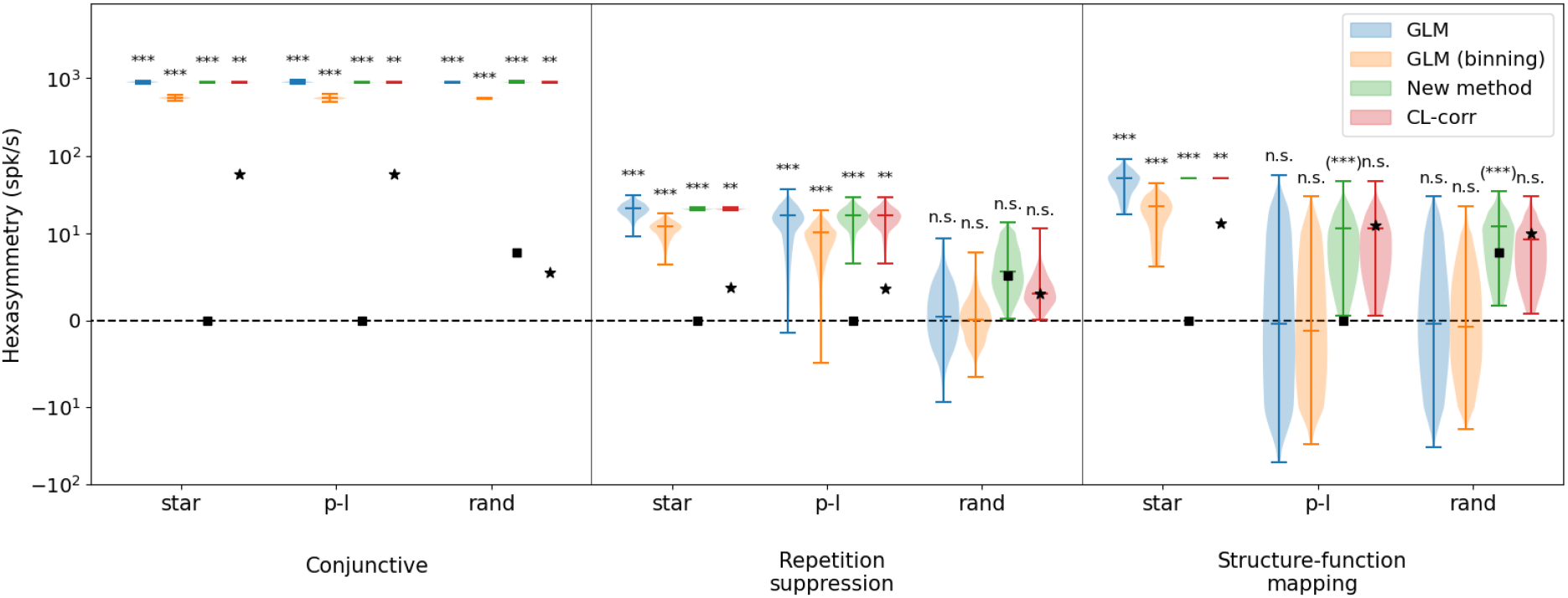
Comparison of hexasymmetry resulting from the three hypotheses when using different methods to calculate the hexasymmetry. Analogous to Fig. 5, each of the three hypotheses (“conjunctive”, “repetition suppression”, “structure-function mapping”) is implemented for three different types of trajectories: star-like walks (“star”), piecewise linear walks (“p-l”), and random walks with small step size (“rand”). For each trajectory type within each hypothesis condition, we show four violin plots with the color corresponding to hexasymmetry calculated using different methods: a generalized linear model (“GLM”), a generalized linear model with binning (“GLM (binning)”), our new method (“New method”), and a circular-linear correlation (“CL-corr”). The mean path hexasymmetry is indicated by a black square. The black star marks the mean hexasymmetry of a surrogate distribution obtained by circularly shifting the firing rates. In all cases, “ideal” parameters were used (conjunctive: *p*_*c*_ = 1, *κ*_*c*_ = 50 rad^*−*2^, *σ*_*c*_ = 0; repetition suppression: *τ*_*r*_ = 3 s, *w*_*r*_ = 1; clustering: *κ*_*s*_ = 10). In order to visualize the GLM methods, which can take on negative values, we use the ‘symlog’ scale provided by pyplot, with a linear threshold of *±* 10 spk/s. Each hypothesis condition for each method was simulated for 300 realizations. ***, *P <* 0.001; **, *P <* 0.01; n.s., not significant.

**Figure S9:**
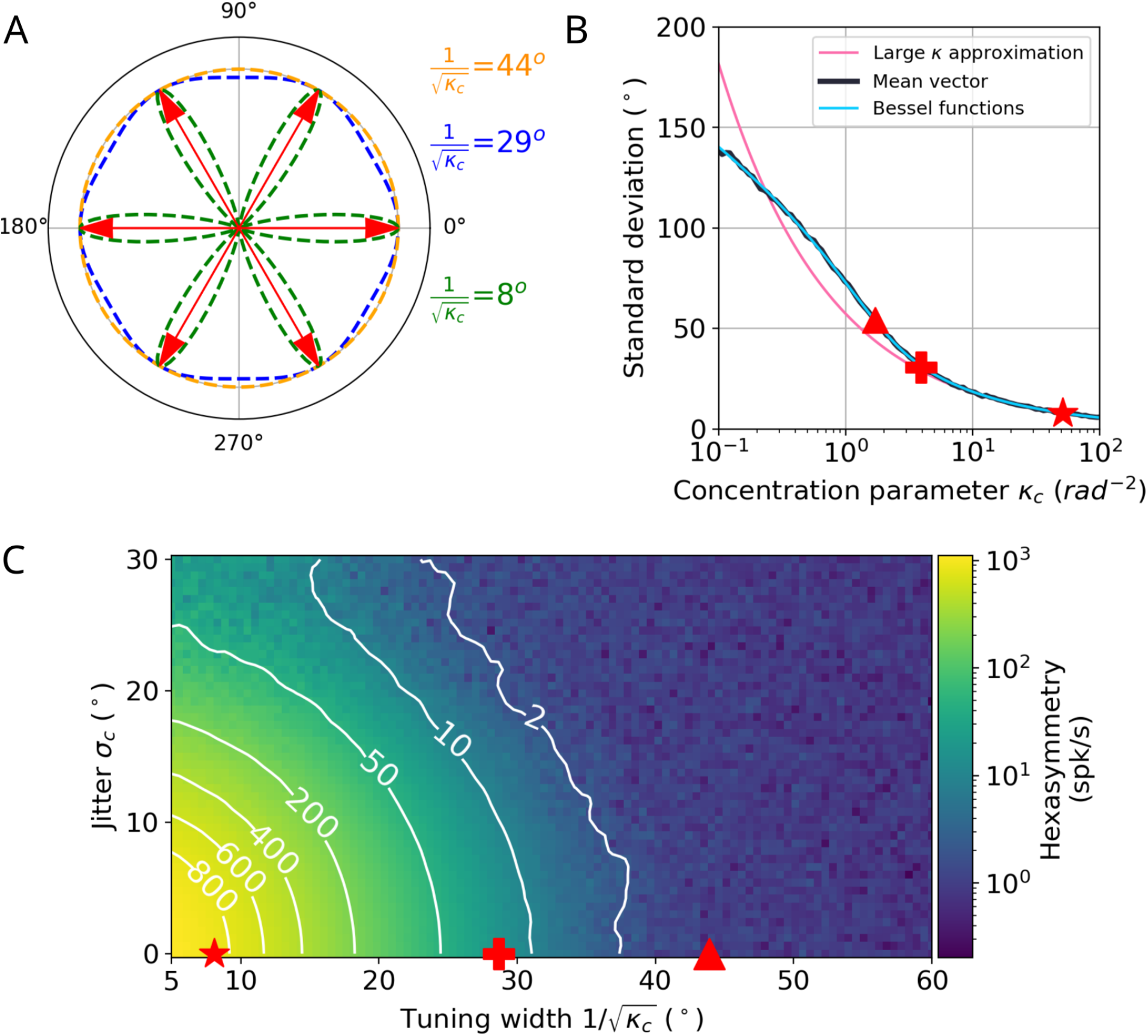
Comparison between the “ideal” tuning width and “realistic” tuning widths found in existing literature for the conjunctive grid by head-direction cell hypothesis. (A) Illustration of tuning curves for different tuning widths when using the large *κ*_*c*_ approximation. (B) Numerical comparison between our tuning widths using the large *κ*_*c*_ approximation and the tuning widths from Sargolini et al. (2006), which use the mean vector approach. The star corresponds to an angular standard deviation of 8.1° (*κ*_*c*_ = 50), which is the tuning width we use for the ‘ideal’ case in the text. The plus denotes an angular standard deviation of 29° (*κ*_*c*_ = 4), which is the tuning width we use for the ‘realistic’ case in the text. The triangle marks an angular standard deviation of 55° (*κ*_*c*_ = 1.7), which is in the range of tuning widths reported by Sargolini et al. (2006). The value of *κ*_*c*_ in this case would be slightly smaller for the same tuning width if the large *κ*_*c*_ approximation is used instead of the mean vector method. “Mean vector” in the legend indicates the angular standard deviation calculated using the vector summation of the mean firing rate over angular bins of 6°. “Bessel functions” denotes the angular standard deviation calculated by using a ratio of Bessel functions instead of the resultant vector. See section 5.8 for details on the calculation of the angular standard deviation. (C) Median hexasymmetry (color coded) as a function of HD tuning width (using the large *κ*_*c*_ approximation) and alignment jitter for star-like walk trajectories. Higher hexasymmetry values are achieved for stronger HD tuning and tighter alignment of the preferred head directions to the grid axes. The star corresponds to a tuning width of 8.1° (*κ*_*c*_ = 50). The plus represents a tuning width of 29° (*κ*_*c*_ = 4). The triangle marks a Sargolini-like tuning width of 44° (*κ*_*c*_ = 1.7) which corresponds to the triangle in panel B. To transform the tuning width from 55° when using the mean vector method to a tuning width of 44° when using the large *κ*_*c*_ approximation, the value of *κ*_*c*_ for a tuning width of 55° was found numerically using the “mean vector” curve in panel B. This value of *κ*_*c*_ was then used in the expression 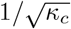 to calculate the tuning width in the large *κ*_*c*_ approximation. The units of 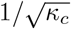 have been transformed from radians to degrees by multiplication with a factor 180*/π*. The hexasymmetry was taken as a median over 4 trials.

**Figure S10:**
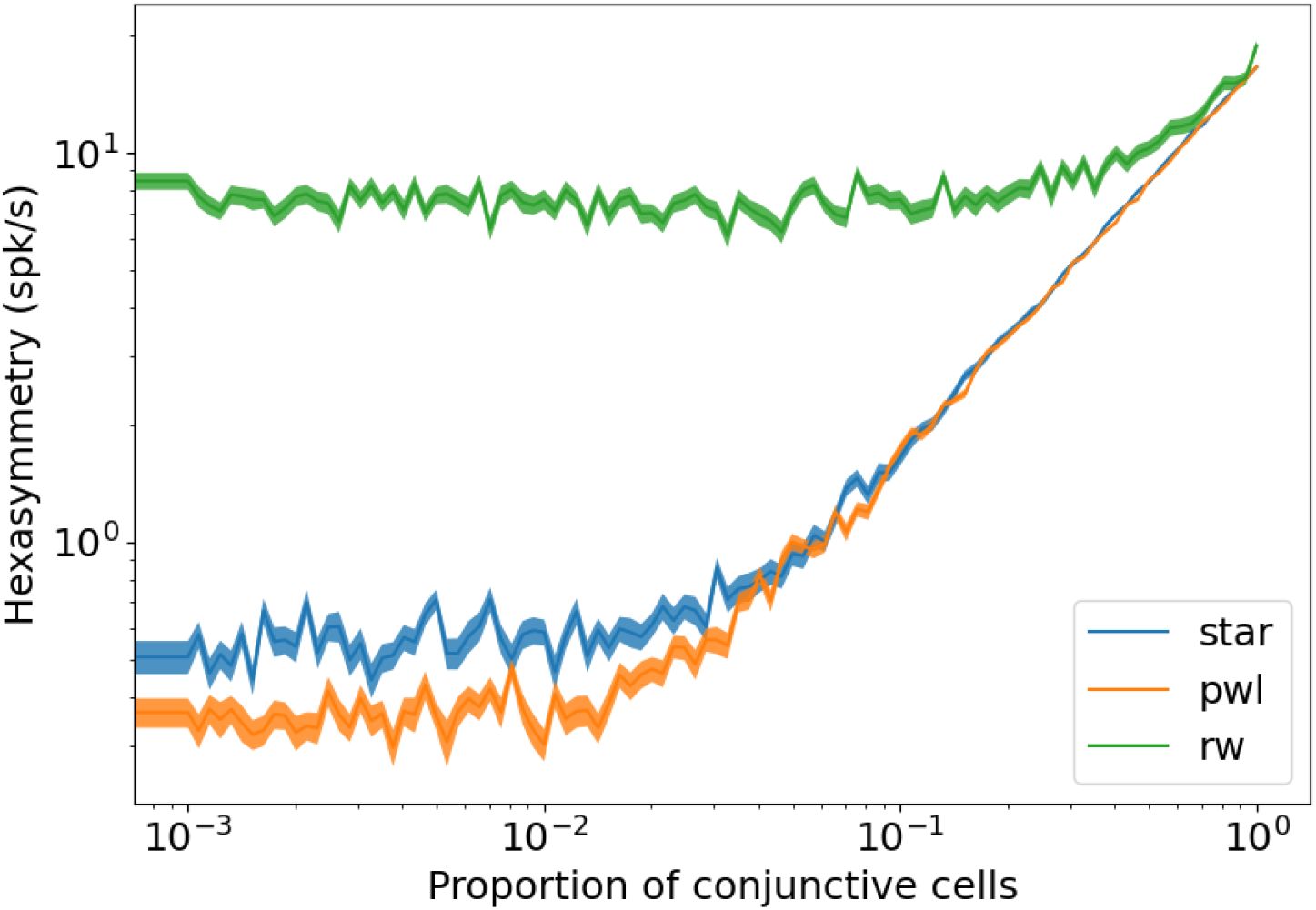
Dependence of the hexasymmetry on the proportion of conjunctive grid by head-direction cells. The hexasymmetry exhibits a linear dependence on the proportion of conjunctive cells up to a noise floor. In random walks, the noise floor is the contribution of the path hexasymmetry to the neural hexasymmetry. For star-like walks and piecewise linear walks, the noise floor is due to sampling a finite number of grid phase offsets from a uniform distribution on the unit rhombus. The “realistic” parameters was used for each trajectory (*p*_*c*_ = 0.33, *κ*_*c*_ = 4 rad^*−*2^, *σ*_*c*_ = 3°). The shaded area represents the standard error of the hexasymmetry

**Figure S11:**
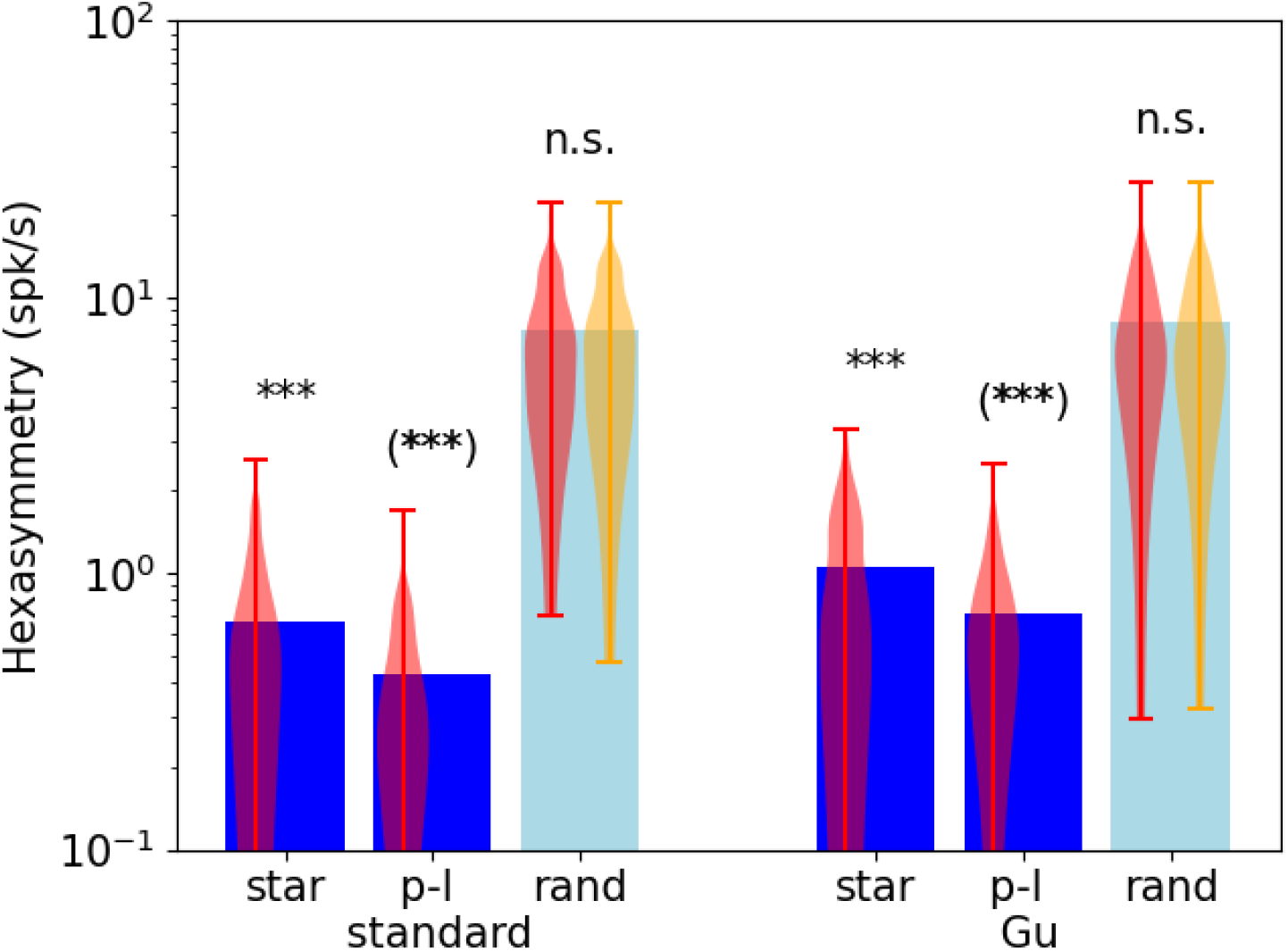
Comparison of hexasymmetry resulting from the standard grid cell model (“standard”) and the structure-function mapping hypothesis (“Gu”, identical to Fig. 5). Each hypothesis is implemented for three different types of trajectories: star-like walks (“star”), piecewise linear random walks (“p-l”), and random walks with small step size (“rand”). For each setting, we show the average neural hexasymmetry |*Ã*_6_ |for 1024 cells (dark blue bars) and the average contribution of the trajectory 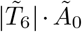 (light blue bars) where *Ã*_0_ is the (variable) average population activity and 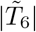 is the hexasymmetry of the trajectory; see Methods for definitions of symbols. The violin plots depict the distributions for firing-rate hexasymmetry (red) and path hexasymmetry (orange). For the “standard” grid cell model, we implemented the structure-function mapping hypothesis, but each realization of the the grid phase offsets was sampled from a two-dimensional von Mises distribution with a clustering parameter *κ*_*s*_ = 0 (equivalent to a uniform distribution on the unit rhombus). In this scenario, random fluctuations give rise to a mean clustering parameter of *κ*_*s*_ = 0.04. The mean clustering parameter in this case depends on the number of grid cells in a voxel (Fig. 4R). Distributing the grid phases regularly throughout the unit rhombus results in *κ*_*s*_ = 0 and reduces the mean neural hexasymmetry (star: 5.6·10^*−*11^ spk/s; p-l: 4.8·10^*−*11^ spk/s). Using a constant function (such that each grid cell has the same firing rate for any position of the agent in space) in place of the grid-like firing fields (Eq. 2) while sampling the grid phase offsets from a uniform distribution on the unit rhombus similarly reduces the mean neural hexasymmetry (star: 3.6 · 10^*−*11^ spk/s; p-l: 3.4 · 10^*−*11^ spk/s). Either regularly distributing the grid phase offsets or using a constant grid function results in a hexasymmetry that is similar in magnitude to the path hexasymmetry. Each hypothesis condition was simulated for 300 realizations. For “Gu”‘ we used *κ*_*s*_ = 0.1. ***, *P <* 0.001; n.s., not significant.

